# Limits in the detection of m^6^A changes using MeRIP/m^6^A-seq

**DOI:** 10.1101/657130

**Authors:** Alexa B.R. McIntyre, Nandan S. Gokhale, Leandro Cerchietti, Samie R. Jaffrey, Stacy M. Horner, Christopher E. Mason

## Abstract

Many cellular mRNAs contain the modified base m^6^A, and recent studies have suggested that various stimuli can lead to changes in m^6^A. The most common method to map m^6^A and to predict changes in m^6^A between conditions is methylated RNA immunoprecipitation sequencing (MeRIP-seq), through which methylated regions are detected as peaks in transcript coverage from immunoprecipitated RNA relative to input RNA. Here, we generated replicate controls and reanalyzed published MeRIP-seq data to estimate reproducibility across experiments. We found that m^6^A peak overlap in mRNAs varies from ∼30 to 60% between studies, even in the same cell type. We then assessed statistical methods to detect changes in m^6^A peaks as distinct from changes in gene expression. However, from these published data sets, we detected few changes under most conditions and were unable to detect consistent changes across studies of similar stimuli. Overall, our work identifies limits to MeRIP-seq reproducibility in the detection both of peaks and of peak changes and proposes improved approaches for analysis of peak changes.

## Introduction

Methylation at the *N*6 position in adenosine (m^6^A) is the most common internal modification in eukaryotic mRNA. A methyltransferase complex composed of METTL3, METTL14, WTAP, VIRMA, and other cofactors catalyzes methylation at DRACH/DRAC motifs, primarily in the last exon (1,2). Most m^6^A methylation occurs during transcription (3). The modification then affects mRNA metabolism through recognition by RNA-binding proteins that regulate processes including translation and mRNA degradation (4–9). However, whether m^6^A is lost and gained in response to various cellular changes remains contentious (3,10–15). To assess the evidence for proposed dynamic changes in m^6^A, a reliable and reproducible method to detect changes in methylation as distinct from changes in gene expression is necessary.

The first and most widely-used method to enable transcriptome-wide studies of m^6^A, MeRIP-seq or m^6^A-seq, involves the immunoprecipitation of m^6^A-modified RNA fragments followed by peak detection through comparison to background gene coverage (16,17). A second method was developed in 2015, miCLIP or m^6^A-CLIP, which involves crosslinking at the site of antibody binding to induce mutations during reverse transcription for single-nucleotide detection of methylated bases (2,18). MeRIP-seq is still more often used than miCLIP, despite less precise localization of m^6^A to peak regions of approximately 50-200 base pairs that can contain multiple DRAC motifs, since it follows a simpler protocol, requires less starting material, and generally produces higher coverage of more transcripts. Antibodies for m^6^A can also detect a second base modification, *N*^6^,2′-*O*-dimethyladenosine (m^6^A_m_), found at a lower abundance than m^6^A and located at the 5′ ends of select transcripts (15,18). We thus refer to the base modifications detected through MeRIP-seq collectively as m6A_(m)_, although most are likely m^6^A. As of late 2018, over fifty studies used MeRIP-seq to detect m^6^A_(m)_ in mammalian mRNA (**Additional File 1: Supplementary Table 1**).

**Table 1:**
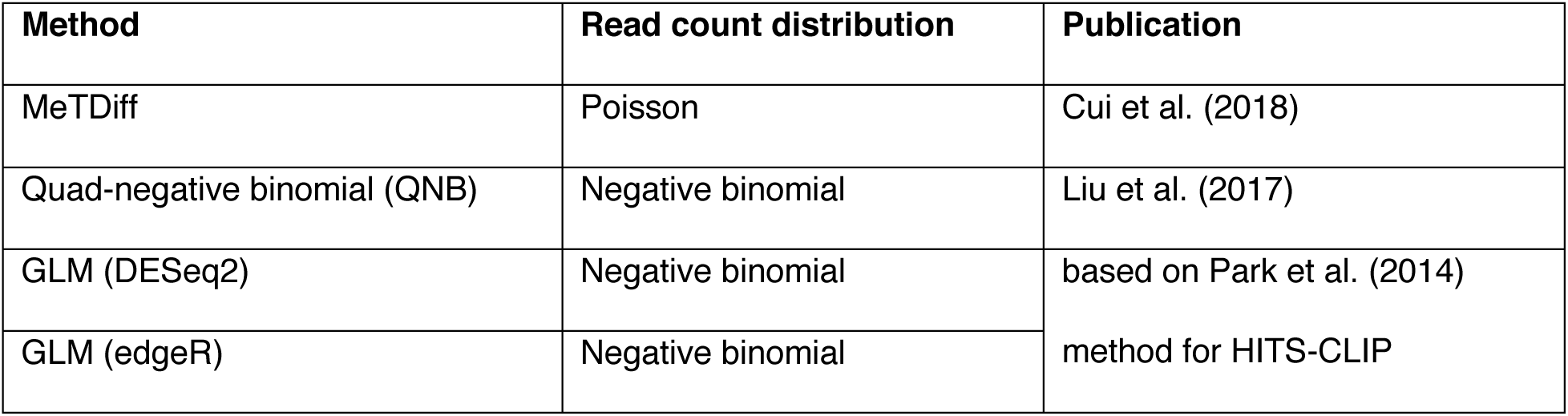
Statistical methods for the detection of peak changes.

Although MeRIP-seq can reveal approximate sites of m^6^A_(m)_, it cannot be used to quantitatively measure the fraction of transcript copies that are methylated (19). Studies of m^6^A variation in response to stimuli instead estimate differences at individual loci through changes in peak presence or peak height.Using these approaches, studies have reported changes to m^6^A with heat shock, microRNA expression, transcription factor expression, cancer, oxidative stress, human immunodeficiency virus (HIV) infection, Kaposi’s sarcoma herpesvirus (KSHV) infection, and Zika virus infection, including hundreds to thousands of changes in enrichment at specific sites (20–29). Statistical approaches to analysis have only recently been published and there have been no comprehensive evaluations of methods to detect changes in m^6^A based on MeRIP-seq data (30,31). Thus, while studies have suggested that m^6^A shows widespread changes in response to diverse stimuli, they have applied inconsistent analysis methods to detect changes in m^6^A and often don’t control for differences in RNA expression between conditions or typical variability in peak heights between replicates. In some cases, these studies have reported m^6^A changes based on simple differences in peak count (24,26,27,32). However, others have applied statistical tests or thresholds for differences in immunoprecipitated (IP) over input fraction enrichment and visual analysis of coverage plots, and have reported fewer m^6^A changes or suggested that m^6^A is a relatively stable mark (33,34). As in RNA-seq, there is noise in MeRIP-seq, and multiple replicates are therefore necessary to estimate variance and statistically identify the effects of experimental intervention (35–37). To date, only one MeRIP-seq study has used more than three replicates per condition (34), while ten have used only one (17,20,32,33,38–43), suggesting that most studies may not have enough power to detect changes in m^6^A_(m)_.

To re-evaluate the evidence for m^6^A_(m)_ changes under various conditions, we first examined the variability in m^6^A_(m)_ detection across replicates, cell lines, and experiments using our own negative controls (12 replicates) as well as 24 published MeRIP-seq data sets. We then compared statistical methods to detect differences in IP enrichment using biological negative and positive controls for m^6^A changes. We found that these methods are limited by noise, including biological variability from changes in RNA expression and technical variability from immunoprecipitation and sequencing that limits reproducibility across studies. Our results suggest that the scale of statistically detectable m^6^A_(m)_ changes in response to various stimuli is orders of magnitude lower than the scale of changes reported in many studies. However, we also found that statistical detection could miss the majority of changed sites when using only 2-3 replicates. We use our results to propose approaches to MeRIP-seq experimental design and analysis to improve reproducibility and more accurately measure differential regulation of m6A_(m)_ in response to stimuli. These data and analyses emphasize the need for further research and alternative assays, for example recently developed endoribonuclease-based sequencing methods (44,45) or direct RNA nanopore sequencing (46), to resolve the extent to which m^6^A changes in response to specific conditions.

## Results

### Detection of peaks across replicates, experiments, and cell types

The first steps in MeRIP-seq data analysis are to align sequencing reads to the genome or transcriptome of origin and to identify peaks in transcript coverage in the IP fraction relative to the input control. Several methods have been developed for MeRIP-seq peak detection, including exomePeak, MeTPeak, MeTDiff, and bespoke scripts. Another method often used for MeRIP-seq peak detection is MACS2, which was originally designed to detect protein binding sites in DNA from chromatin immunoprecipitation sequencing (ChIP-seq). We compared m^6^A_(m)_ peak detection by exomePeak, MeTPeak, MeTDiff, and MACS2 (31,47–49) in seven replicates of MeRIP-seq data obtained from mouse cortices under basal conditions (34), and in 12 replicates of MeRIP-seq data we generated from human liver Huh7 cells (50). The intersect between all tools tested was high, and we saw minimal differences in DRAC motif enrichment, which we use to provide an estimate of tool precision in the absence of true positive m^6^A sites (**Additional File 2: Supplementary Figure 1a**). In addition, we assessed the METTL3/METTL14-dependence of specific peaks identified by single tools using MeRIP-RT-qPCR. We found that of these peaks, 4/4 from MACS2, 5/5 from MeTPeak, and 4/5 from MeTDiff showed decreased m^6^A_(m)_ enrichment following METTL3/METTL14 depletion, suggesting that these are true m^6^A sites. By comparison, only 1/5 of the peaks uniquely called by exomePeak showed statistically significant decreases (p < 0.05), although replicate variance was high and 4/5 showed a downward trend (**Additional File 2: Supplementary Figure 1b**). Since MACS2 was the most commonly used tool for peak calling and was previously found to perform well in comparison with a graphical user interface tool and several other peak callers (51), we used MACS2 for the remainder of our analyses. Repeating the analyses shown in Figures 2-4 using the MeTDiff peak caller instead of MACS2 did not affect any of our conclusions (**Additional File 3**).

**Figure 1:**
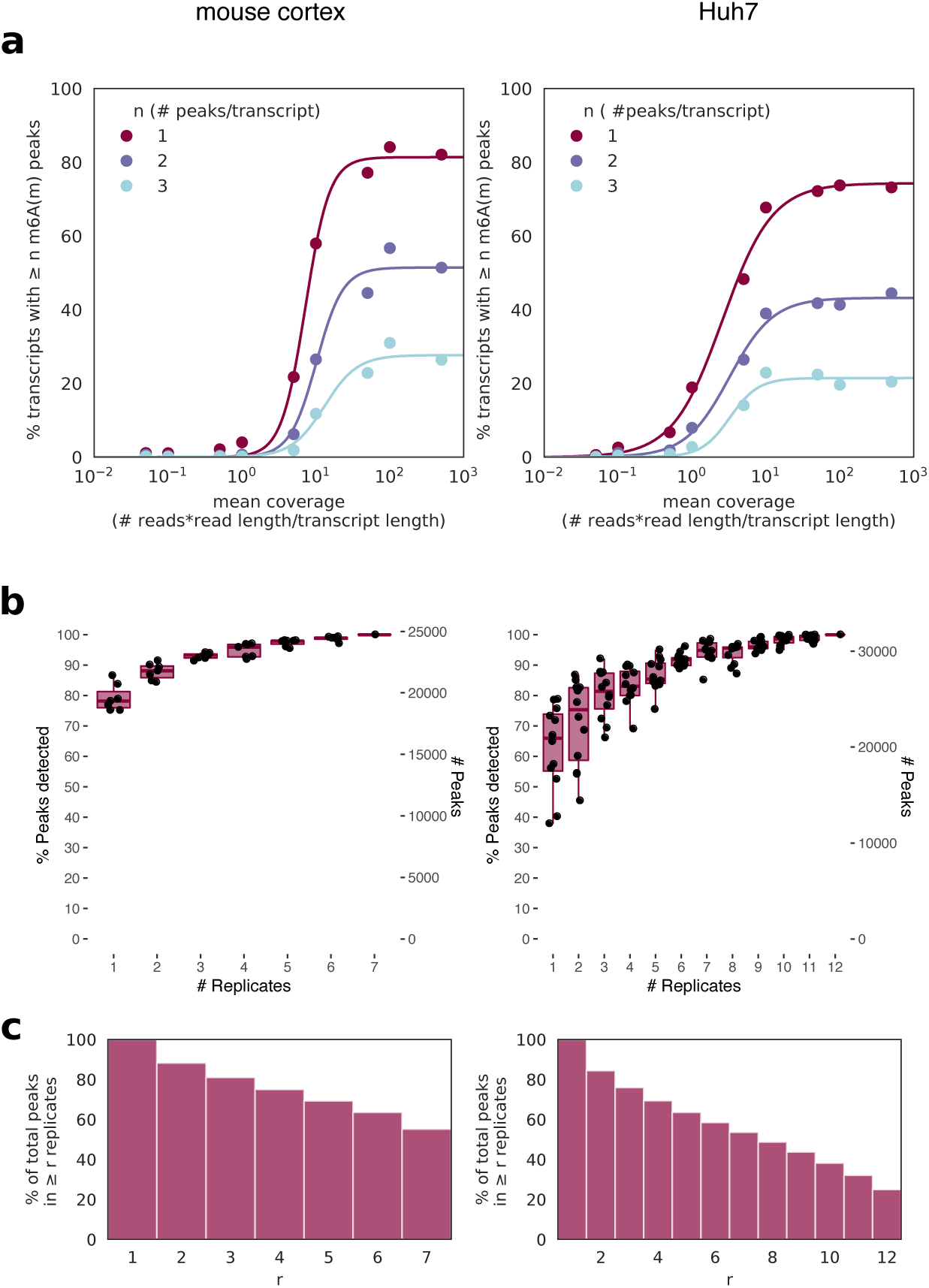
Thresholds for peak detection. **a)** m^6^A_(m)_ site detection in MeRIP-seq data from mouse cortex (left) and human liver cells (Huh7, right) shows saturation of peak detection as transcript coverage approaches 10-50X for replicates at basal conditions, with peaks merged from all replicates. **b)** The total number of peaks captured increases with more replicates, with single replicates capturing a median of 66-78% of total peaks depending on study. Boxes span the 1^st^ to 3^rd^ quartiles of distributions for random subsamples of replicates, with lines indicating the median number of peaks, and whiskers showing the minimum and maximum points within ±1.5x the interquartile distance from the boxes. Jittered points show results for each random subsample (a total of 6 subsamples per replicate number for the mouse cortex data and 12 for the Huh7 data). **c)** The percent of peaks detected in at least r replicates for the same data sets.

**Figure 2:**
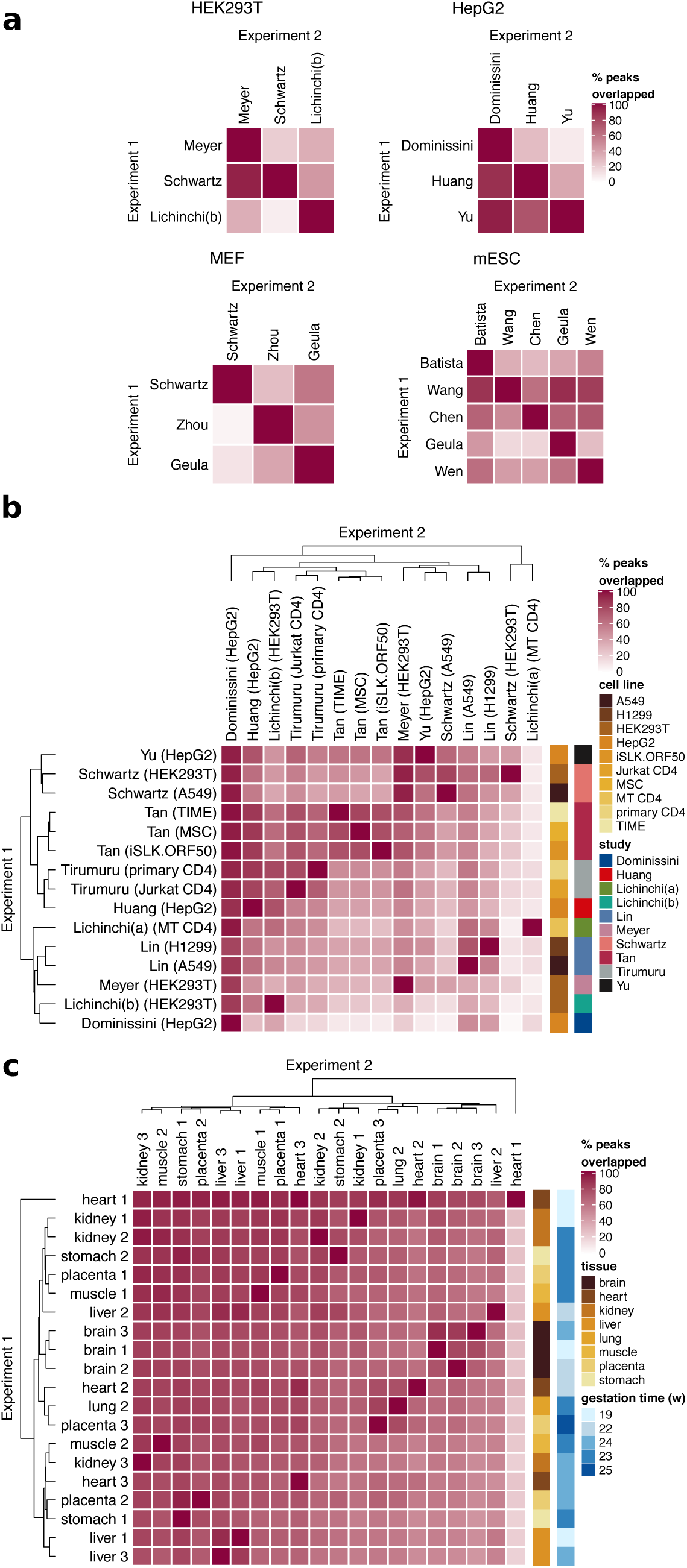
Reproducibility of peak detection. **a)** Peak detection between studies that used the same cell type shows variable overlap. Overlap was calculated as the percent of peaks detected in Experiment 1 with an overlap of ≥ 1 base pair with peaks from Experiment 2. **b)** Peak detection across tissue and cell types shows samples from the same study cluster better together than samples from the same tissue. Median overlap was 46%. **c)** Peak detection across tissue types for data from the same study (Xiao et al., 2019). Median overlap was 72%. Studies used in (a) and (b) are described in **Additional File 1: Supplementary Table 2**

For m^6^A_(m)_ peak detection, a transcript must be sufficiently expressed for enrichment by the m^6^A_(m)_ antibody and for adequate sequencing coverage in both the IP and input fractions. Previous reports have suggested that m^6^A_(m)_ presence does not decrease with lower mRNA expression level, and, if anything, is higher in mRNAs with lower expression as methylated transcripts tend to be less stable (9,38). Peak callers, however, identify fewer peaks in genes at low expression, which we therefore assume reflects inadequate coverage for peak calling. To estimate the level of coverage necessary for peak detection, we analyzed the percent of genes with at least one, two, or three peaks relative to mean input transcript coverage in both the mouse cortex and Huh7 cell data (**Figure 1a**). Based on the upper shoulders of the sigmoidal curves as the percent of genes with peaks begins to plateau, we estimate that mean gene coverage of approximately 10-50X is necessary to avoid missing peaks based on insufficient coverage. Including a wider array of samples in this analysis likewise showed an increase in the percent of transcripts with ≥1 peak as coverage rose to 10X (**Additional File 2: Supplementary Figure 1c**). Our analysis of the input RNA-seq coverage of peak regions alone again supported a similar threshold; few peaks are detected with median input read counts below 10 across replicates (**Additional File 2: Supplementary Figure 1d**). These thresholds do not mean that peaks in genes with mean coverage < 10X or peaks with fewer than 10 input reads are false positives, but that the likelihood of false negatives rises with lower coverage (**Additional File 2: Supplementary Figure 1e**).

To evaluate the reproducibility of MeRIP-seq data, we next examined the consistency of m^6^A_(m)_ peak calling between replicates. Previous studies have reported that peak overlap between replicates is approximately 80% (9,16,52,53). Similarly, we found that between two replicates, log_2_ fold enrichment of IP over input reads at detected peaks showed a Pearson correlation of approximately 0.81 to 0.86 (**Additional File 2: Supplementary Figure 1f**). A single sample captured a median of 78% of the peaks found in seven replicates of mouse cortex data and 66% of peaks found in twelve replicates of Huh7 cell data. The number of detected peaks increased log-linearly with the addition of more replicates, such that with three replicates, 84-92% of the peaks found with 7-12 replicates were detected (**Figure 1b**). Conversely, the number of peaks in common across replicates decreased as the number of replicates increased, such that while ∼ 80% of peaks were detected in at least two replicates, only ∼ 60% were detected in six replicates for both data sets and ∼ 25% in all twelve replicates of Huh7 cell data (**Figure 1c**). Detection of peaks in more replicates did not increase DRAC motif enrichment (**Additional File 2: Supplementary Figure 1g**). These results suggest that many m^6^A_(m)_ sites may be missed in studies that use one to three replicates, and that increasing replicates could enable detection of more peaks.However, not all peaks correspond to true m^6^A_(m)_ sites. A recent comparison to data from an endoribonuclease-based method for m^6^A detection suggested MeRIP-seq has a false positive rate of ∼ 11%, although this would differ by study and detection threshold (3,54).

The number of peaks detected across studies varies. Given that coverage affects peak detection, we hypothesized that variation in sequencing depth could contribute to differences in peak count. Zeng et al. (2018) reported that peak count begins to saturate by around 20 million reads by subsampling data within individual studies (42). However, we found that there is no positive correlation between peak count and input or IP sequencing depth across data sets from different published studies, each of which had 3-81M reads per replicate (input Pearson’s R = −0.37, p = 0.015; IP Pearson’ s R = −0.17, p = 0.28) (**Additional File 1: Supplementary Table 2, Additional File 2: Supplementary Figure 2a-b**). This implies that experimental factors beyond sequencing depth contribute to the variability of peak counts across studies.

We next analyzed the overlap of peaks among studies and found inconsistency in peak localization on transcripts as well. Within four commonly used cell types, the percent of peaks detected in one experiment that were also detected in a second varied among pairs of studies from as low as 2% of peaks to as high as 90% (median = 45%), after filtering for transcripts expressed above a mean of 10X input coverage in both to ensure sufficient expression for peak detection (**Figure 2a**). In fact, peaks showed higher overlap within different cell types from the same study than within the same cell type from different studies, suggesting that MeRIP-seq data is prone to strong batch effects (**Figure 2b**). While this could be due to differences among experimental protocols used (summarized in **Additional File 1: Supplementary Table 2**), we were unable to identify such a link. Overall, most percent overlaps of m^6^A_(m)_ peaks fell between ∼30% (1^st^ quartile) and ∼60% (3^rd^ quartile) (**Figure 2b**). With rare exceptions (e.g. that described by Ke et al., 2017 in their Supplementary Figure 8 (3)), most MeRIP-seq data sets do show enrichment of the m^6^A motif DRAC. These results indicate, however, that multiple labs running MeRIP-seq on the same cell type will detect different subsets of m^6^A_(m)_ sites. Possible contributing factors in the differences among studies include cell state (e.g. different stages of the cell cycle), experimental conditions, and sequencing depth. Despite predictions that tissue or cell type would be a large factor in differences among samples, though, peaks detected in different tissues analyzed in a single experiment showed high overlap and little clustering by tissue type (**Figure 2c**) (55). This suggests that although there is evidence that m^6^A levels vary by tissue (19), modified sites are consistent.

### Detection of changes in peaks between conditions

Following m^6^A_(m)_ peak detection, many studies compare the expression of peaks between two conditions to predict peak changes. While looking at plots of IP and input gene coverage under different conditions can help evaluate the evidence for these changes (33), statistical or heuristic methods are first necessary to narrow down a list of candidate sites to plot. Several tools used for statistical analysis by the studies in **Additional File 1: Supplementary Table 1** or for other types of RNA IP sequencing assays model peak counts using either (a) the Poisson distribution, in which the variance of a measure (here, read counts) is assumed to be equal to the mean (MeTDiff), or (b) the negative binomial distribution, in which a second parameter allows for independent adjustment of mean and variance (QNB and two implementations of a generalized linear model approach using DESeq2 or edgeR, **Table 1**) (30,31,56– 58). In the mouse cortex and Huh7 cell data, we found that, similar to RNA-seq data (24,57,59), the variance in read counts under peaks exceeded their mean, indicative of overdispersion (**Additional File 2: Supplementary Figure 3a**). The log likelihood (the probability of an observation given a distribution with known parameters) for our sample also fell within the distribution of expected log likelihoods for the negative binomial distribution (bottom) but not the Poisson distribution (top) (**Figure 3a**). Thus, the negative binomial distribution captures the mean-variance relationship in MeRIP-seq data, suggesting that tools that account for overdispersion better model the distribution of read counts at m^6^A_(m)_ peaks than tools that do not.

**Figure 3:**
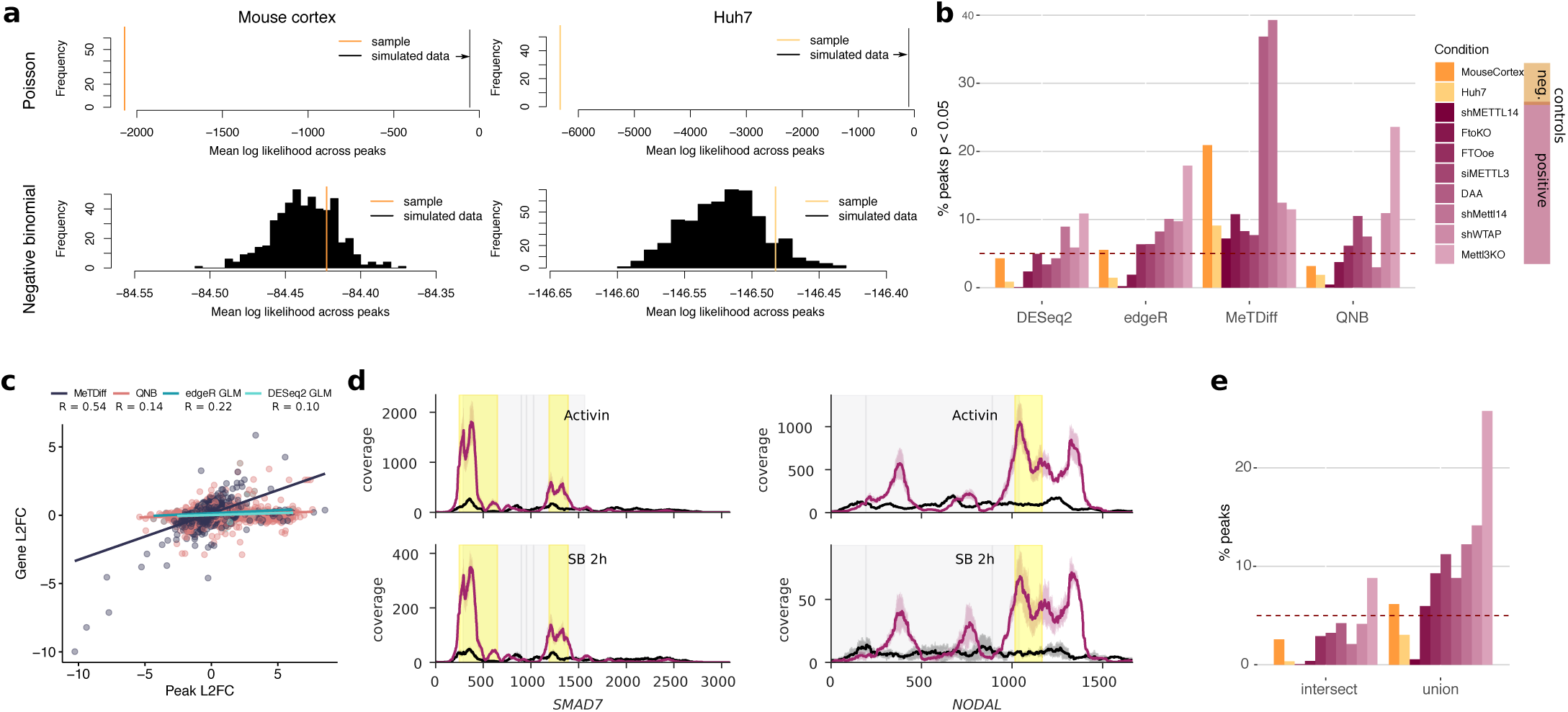
Analysis of methods to detect peak changes disproportional to gene expression changes. **a)** A comparison of Poisson (above) and negative binomial (below) models for read counts under peaks. The negative binomial mean log likelihood of the sample data fell within the 74^th^ and 89^st^ percentiles of 500 simulations for mouse cortex and Huh7 cell data, respectively, while the Poisson model failed to capture the sample distributions. **b)** The percent of sites below an unadjusted p-value threshold of 0.05 for different methods (described in **Table 1**) to detect differential methylation in negative controls between two groups at baseline conditions and positive controls in which methylation processes were disrupted with respect to baseline conditions (**Additional File 1: Supplementary Table 3**). The line at 5% indicates the expected proportion of sites given a uniform p-value distribution (see **Additional File 2: Supplementary Figure 2c**), while colours indicate negative (orange) and positive (purple) control experiments. **c)** The correlation between change in gene expression and change in peak expression between conditions for sites identified as differentially methylated in the eight positive control experiments. Pearson’s R = 0.22, 0.10, 0.55, and 0.14 for edgeR, DESeq2, MeTDiff, and QNB, respectively, with p = 0.05, 0.09, 5.8E-87, and 2.4E-11. **d)** Coverage plots showing changes in peak expression are proportional to changes in gene expression for genes identified as differentially methylated by Bertero et al. (2018) using MeTDiff with activin signaling and an activin-NODAL inhibitor, SB431542 (SB). Lines show the mean coverage across three replicates, while shading shows the standard deviation. Peaks detected as significantly changed are highlighted in yellow. Coding sequences are shown in grey. **e)** The intersect and union of peaks with p < 0.05 from DESeq2, edgeR, and QNB from (b), coloured as in (b).

We next defined positive and negative controls to evaluate tool performance for detection of changes in m^6^A_(m)_ peaks. Past publications describing new methods to detect m^6^A_(m)_ peak changes have used data sets in which methylation machinery genes or the methyl donor were disrupted compared to baseline conditions as positive controls, and have simulated negative controls by randomly swapping labels in the positive controls (30,31). However, swapping labels for conditions that may feature differences in gene expression in addition to differences in m^6^A levels could unrealistically increase variance in read counts within groups. Therefore, we instead used the two data sets from mouse cortex and Huh7 cells, which each comprised many replicates at baseline conditions (n=7 and n=12, respectively), as negative controls. We randomly divided the mouse cortex data into two groups of three to four replicates for comparison and divided the Huh7 replicates by lab of incubation, which did not affect sample clustering (**Additional File 2: Supplementary Figure 3b**). We would expect to see minimal changes in IP enrichment at m^6^A peaks between groups for our negative controls, whereas our positive controls, which featured genetic or chemical interference with the m^6^A machinery, should show discernible differences in peaks when compared to baseline or wildtype conditions in the same cell lines (summarized in **Additional File 1: Supplementary Table 3**). Indeed, the absolute difference in log_2_ fold change between peaks and genes was centered around 0 for the negative controls and showed small shifts that varied in magnitude and direction for the positive controls (**Additional File 2: Supplementary Figure 3c**).

Using statistical methods to detect changes in peak enrichment, we found that the percent of changes called below a p-value threshold of 0.05 were similar in the positive and negative controls (**Figure 3b**). With all tools except MeTDiff, a knockout of Mettl3 showed the largest effects on m^6^A (60), while fewer significant peaks in other positive controls suggested variable effects of the positive control conditions on m^6^A_(m)_, possibly related to efficiency of the methylation machinery knockdown or overexpression (7,33,61–65). In the absence of true differences between groups, p-value distributions should be uniform for well-calibrated statistical tests, meaning that ∼5% of peaks should have p-values < 0.05 for the negative controls. MeTDiff reported an excess number of sites with p-values below 0.05 (**Additional File 2: Supplementary Figure 3d**) and identified a higher percentage of sites as differentially methylated in the mouse cortex negative control data set than in all but two positive controls (**Figure 3b**). On the other hand, the generalized linear models (GLMs) and QNB showed uniform to conservatively shifted p-value distributions, with differences between the mouse cortex and Huh7 data sets (**Additional File 2: Supplementary Figure 3d**), suggesting that these tools detect fewer false positives.

To ensure significant peak changes detected by each of the tools reflected changes in IP enrichment independent of differential gene expression, we measured the correlation between changes in IP read counts at peak sites and changes in input read counts across their encompassing genes. For significant peaks (FDR-adjusted p-value < 0.05) from the positive controls, correlation between log_2_ fold change in peak IP and gene input read counts was low for the GLMs and QNB (Pearson’s R = 0.10 to 0.22) but reached 0.55 (p = 5.8E-87) for MeTDiff (**Figure 3c**). The higher correlation for MeTDiff was driven by peaks with proportional changes in IP and input levels, which suggests that MeTDiff often detects differential expression of methylated genes rather than differential methylation. Therefore, published studies that have used MeTDiff may actually be detecting differential expression and not differential methylation (22,66). Indeed, plotting coverage for genes reported as differentially methylated in one of these studies, with the y-axis scaled separately per condition, confirmed that changes in m^6^A identified by MeTDiff were proportional to changes in gene expression (**Figure 3d**) (22). Given these results, QNB or the GLM implementations are better methods than MeTDiff to detect differential methylation. Taking the intersect of significant peaks for the GLMs and QNB may help determine the most probable sites of m^6^A changes, while taking the union of predictions provides a less conservative approach to selecting sites for further validation (**Figure 3e**). However, additional filters are needed for robust peak change detection as there were still significant peaks for which the difference between peak log_2_ fold change and gene log_2_ fold change was close to zero, particularly with QNB (**Additional File 2: Supplementary Figure 3e**). For microarray and RNA-seq data, a filter of absolute log_2_ fold change > 1 has been recommended to reduce false positive rates (67); in the remainder of our analyses, we implemented a similar filter for absolute difference in peak and gene log_2_ fold change ≥ 1 to the combined predictions from QNB and the two GLMs, with an additional filter where noted for peak read counts ≥ 10 across all replicates and conditions to ensure sufficient coverage for consistent peak detection (as discussed in Figure 1a).

### Reanalyzing peak changes between conditions

We next estimated the scale of statistically detectable peak changes under various conditions using our approaches and compared these results to previously reported estimates of these changes (**Figure 4a, Additional File 1: Supplementary Table 4**). We identified fewer peaks as differentially methylated than originally reported under most conditions, with zero to hundreds of peaks significantly changed (depending on experiment and method), versus hundreds to over ten thousand described in publications (22–26,34,63,66,68–71). Notably, knockdown of Zc3h13 did appear to disrupt m^6^A_(m)_, suggesting the gene does participate in methylation as recently described (69). Another study reported that activin treatment of human pluripotent stem cells led to differential methylation of genes that encode pluripotency factors (22). However, our reanalysis only found a few peak changes that passed our filters for significance, fold change, and expression (minimum input read count across peaks ≥ 10), and no enrichment for pluripotency factors among affected genes. Even when we removed the thresholds for fold change and expression, the adjusted p-value for “signaling pathways regulating pluripotency of stem cells” was still 0.15 and driven by only three genes, LEFTY2, FZD28, and FGFR3 (**Additional File 2: Supplementary Figure 4a**). Interestingly, the minimum read threshold made a particularly dramatic difference in the case of a recent study that looked at the effects of knocking down the histone methyltransferase SETD2 on m^6^A in mRNA. For this data, of the 2065 sites predicted by QNB, 2064 fell below the minimum read threshold due to low input coverage in the first and second replicates (**Figure 4a, Additional File 2: Supplementary Figure 4b-e**) (70). We could not compare our approach to results reported by Su et al. (2018), who found 6,024 peaks changed with R2HG treatment, Zeng et al. (2018), who found 465-599 peaks changed between tumour samples, or Ma et al. (2018), who found 12,452 peaks were gained and 11,192 lost between P7 and P20 mouse cerebella, as each relied on a single sample per condition, with no replicates (40–42).

**Figure 4:**
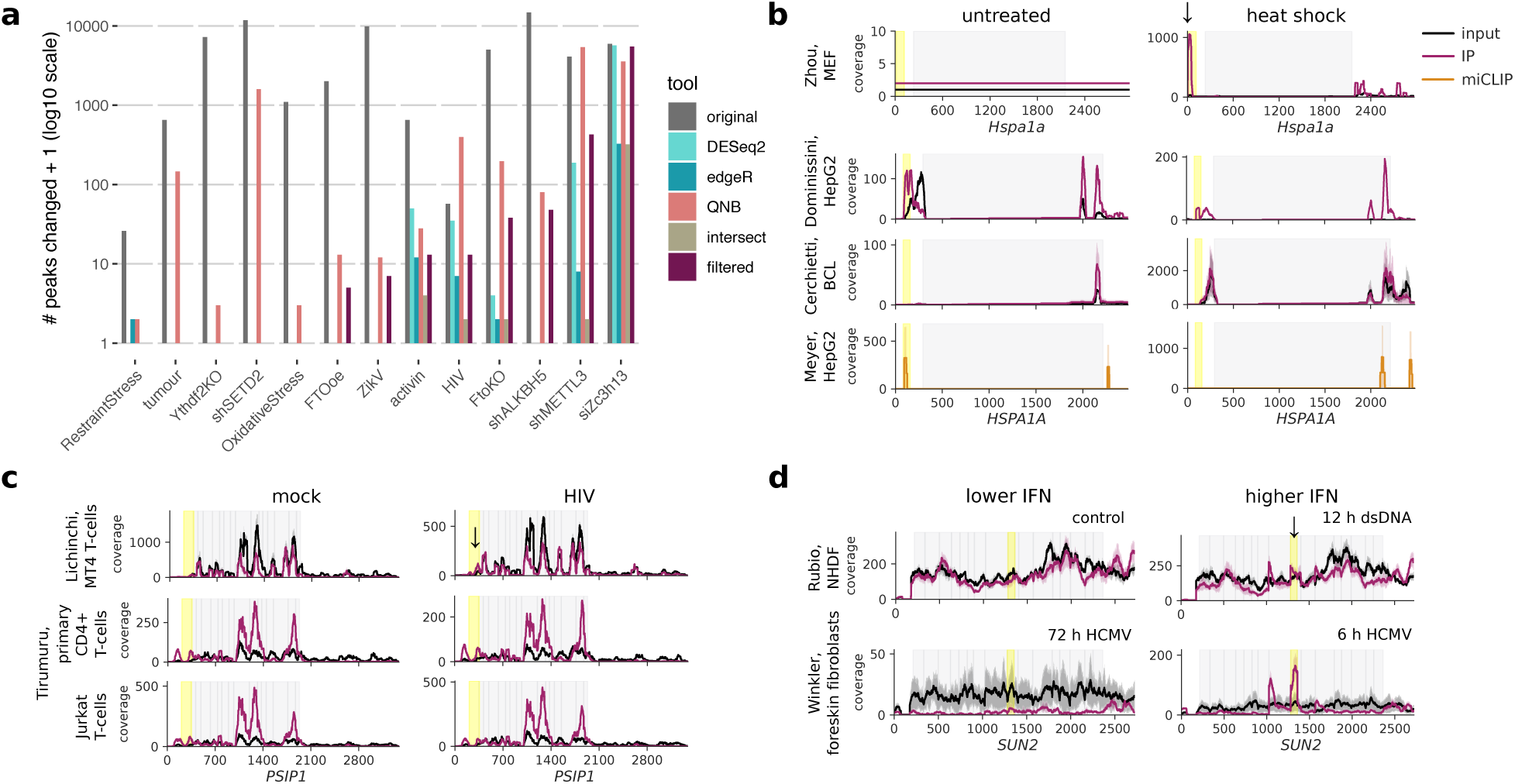
Changes in peaks between conditions. **a)** Detected m^6^A_(m)_ changes in thirteen published data sets that measured m^6^A_(m)_ peak changes between two conditions **(Additional File 1: Supplementary Table 4)**. The number of peaks detected as changed in the original published analyses are compared to the number of peaks with FDR-adjusted p-values < 0.05 in our reanalysis using DESeq2, edgeR, or QNB, and taking the union of results from these three tools with additional filters for log_2_ fold difference in peak and gene changes of ≥1 and peak read counts ≥10 across all replicates and conditions (“filtered”). **b)** Gene coverage plots for *Hspa1a* in mouse embryonic fibroblasts (MEFs) and *HSPA1A* in human cells (HepG2 and BCL) before and after heat shock. Input coverage is shown in black and IP coverage in raspberry, with putative m^6^A peaks changed highlighted in yellow and marked by arrows. miCLIP coverage for an experiment in HepG2 cells is shown in orange.**c)** Coverage plots for *PSIP1*, which was reported to have a change in 5′ UTR m^6^A with HIV infection by Lichinchi et al (2016). **d)** Coverage plots for *SUN2*, in which we detected changes in m^6^A with HCMV infection and dsDNA treatment suggesting a possible increase in methylation under higher interferon conditions (after 12 h of dsDNA treatment compared to untreated controls and after 6 h post-HCMV infection compared to 72 h, when interferon levels have declined). Lines in coverage plots (**b-d**) show the mean across all replicates for each experiment, while shading shows the standard deviation. Coding sequences are shown in grey.

Multiple studies have investigated m^6^A_(m)_ in the context of heat shock, HIV infection, KSHV infection, and dsDNA treatment or human cytomegalovirus (HCMV) infection (**Additional File 1: Supplementary Table 5**). Since each step in MeRIP-seq analysis risks introducing false negatives, we cannot rule out consistent changes between studies that used similar experimental interventions based on statistical detection alone. Therefore, we started by plotting coverage for specific genes reported as differentially methylated to evaluate reproducibility across these studies. Zhou, et al. (2015) reported 5 ′ UTR methylation of *Hspa1a* with heat shock (20). Coverage was too low for untreated controls to determine if *Hspa1a* was simply newly expressed or was actually newly methylated with heat shock based on our alignment of their data using STAR (72). We were also unable to detect a change in methylation of *HSPA1A* using data from other heat shock studies, including a new data set from a B-cell lymphoma cell line and a published miCLIP data set, although coverage was again low (**Figure 4b**) (4,73). Lichinchi, et al. (2016) reported that 56 genes showed increased methylation with HIV infection in MT4 T-cells, with enrichment for genes involved in viral gene expression (25). Specific genes, for example *PSIP1*, in which we also detected a peak using MACS2 and see a change in the peak when plotting coverage using the data from Lichinchi et al. (2016), did not show the same changes in data from two other CD4^+^ cell types, primary CD4^+^ cells and Jurkat cells (**Figure 4c**) (74). Two other studies both used MeRIP-seq to establish the presence of m^6^A in *IFNB1* induced through dsDNA treatment or by infection with the dsDNA virus HCMV (75,76). While these studies did not discuss changes in m^6^A, we used these data sets to examine the replicability of m^6^A_(m)_ changes in response to dsDNA sensing and interferon induction. Although different dsDNA stimuli, time points, and use of a fibroblast cell line versus primary foreskin fibroblasts make it difficult to compare between the two experiments, using QNB and the GLM approaches, we found four peaks in three genes (*AKAP8, SUN2*, and *TMEM140*) that showed significant changes with higher interferon (**Figure 4d**). Overall, we were unable to detect the same changes in m^6^A_(m)_ across studies of heat shock or HIV, and we detected only a few common changes in the response to dsDNA. However, we do note that cell line-specific differences in m^6^A_(m)_ regulation and differences in experimental protocols could account for some of the variability among these studies.

We did not have MeRIP-seq data for two studies from exactly the same conditions and cell lines to compare, but two studies both used cell lines derived from iSLK to study the effects of KSHV on host m^6^A (27,28). Both suggested that KSHV infection could decrease the number of m^6^A sites in host transcripts. Hesser et al. (2018) found that lytic KSHV infection decreased the number of peaks on host transcripts by >25%; Tan et al. (2018) suggested a loss of 17-59% of peaks in two different cell types, but that m^6^A_(m)_ peak fold enrichment showed better clustering by cell type than by infection status. Neither of these studies discussed specific genes that showed differential methylation with lytic infection. For our comparison of m^6^A_(m)_ peak changes in these data sets, we identified probable changes in peaks based on statistical significance using QNB or the GLMs with log_2_ fold change difference between peaks and genes of ≥1. We detected 80 peak changes in the data from Hesser et al. (2018) and 18 in the data from Tan et al. (2018) but found no peaks that changed in both iSLK data sets with lytic KSHV infection. Applying the same statistical approaches, we were likewise unable to detect any shared peak changes between the studies of HIV infection, and there were insufficient replicates to compare heat shock studies (16,20,25,73,74). Thus, in our reanalysis of m^6^A changes in response to stimuli, we detected only four statistically reproducible peak changes, all in response to dsDNA.

Disparities between experiments were not simply due to significance thresholding or differences in peak detection. Taking the union of peaks called in two experiments for KSHV, HIV, and dsDNA treatment, we found minimal to negative correlations in changes in m^6^A enrichment induced by treatment at the same sites, further showing that changes with similar treatments are not reproducible (**Additional File 2: Supplementary Figure 4e**).

### MeRIP-RT-qPCR validation

Although statistical approaches revealed fewer changes in m^6^A_(m)_ with various stimuli than published estimates, and we were unable to confirm changes in m^6^A_(m)_ methylation of specific genes across studies of similar conditions, many of the studies we looked at do include additional validation of m^6^A_(m)_ changes from MeRIP-seq using MeRIP-RT-qPCR. Recently it was shown that MeRIP-RT-qPCR can capture differences in m^6^A:A ratios at specific sites (34), but it is unknown how MeRIP-RT-qPCR is affected by changes in gene expression. To test this, we ran MeRIP-RT-qPCR on in vitro transcribed RNA oligonucleotides that lacked or contained m^6^A spiked into total RNA extracted from Huh7 cells (**Additional File 1: Supplementary Table 6**). We found that MeRIP-RT-qPCR detected the direction of change in m^6^A levels at different concentrations of spike-in RNAs (**Figure 5a-b**). However, technical variation could also lead to spuriously significant differences. For example, a comparison of m^6^A enrichment between two dilutions (0.1 fmol and 10 fmol) of a 30% methylated spike-in mixture returned a p-value of 0.004 (unpaired Student’s *t*-test).

**Figure 5:**
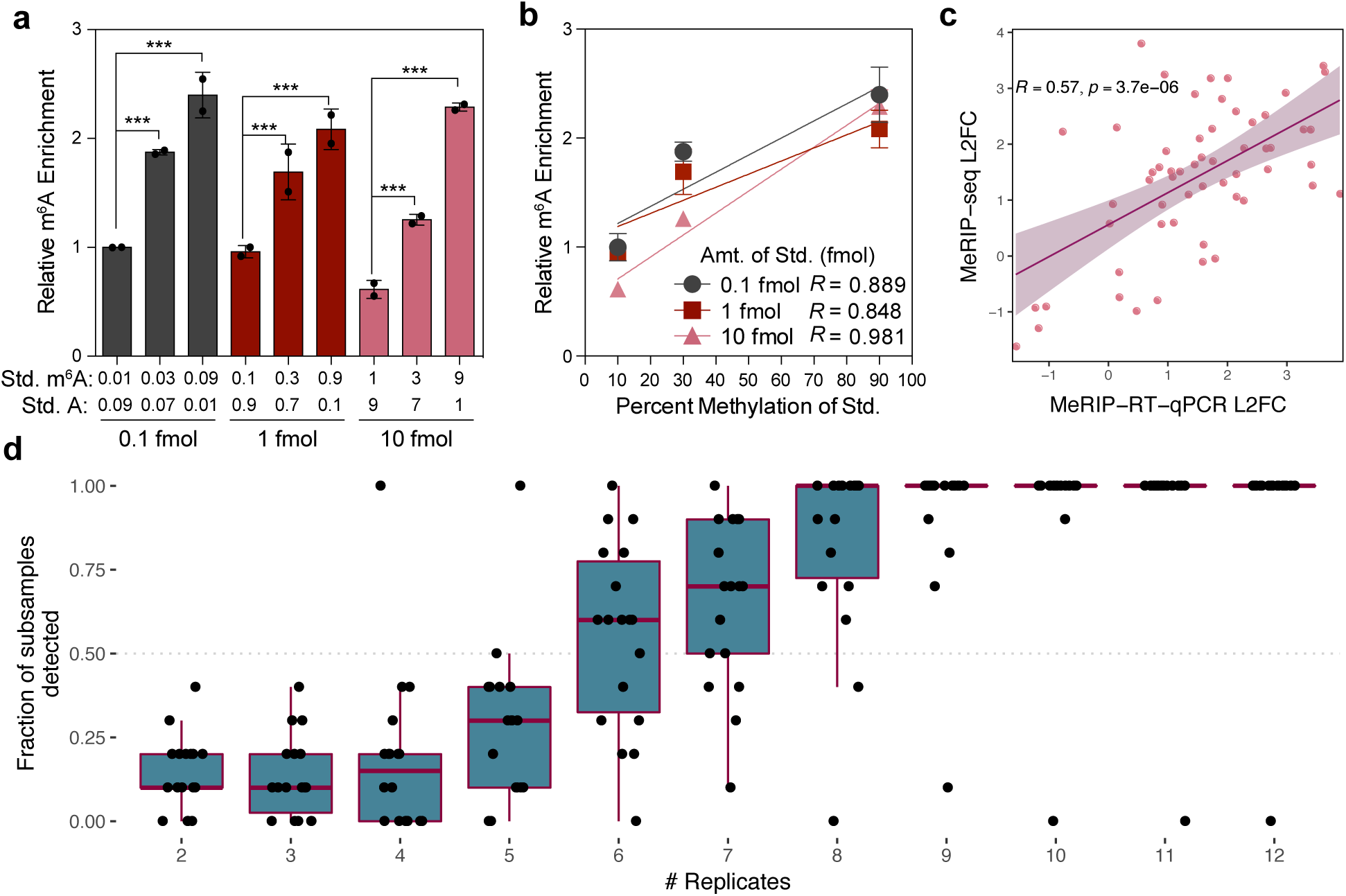
MeRIP-RT-qPCR validation and replicates necessary for the detection of peak changes. **a)** Relative enrichment of the indicated amounts of an in vitro transcribed standard containing unmodified A or m^6^A, as measured by MeRIP-RT-qPCR. Data are shown for two independent replicates of three technical replicates each as IP enrichment over input relative to pulldown of a positive control spike-in, with the 0.1 fmol (0.01 m^6^A: 0.09 A) sample normalized to 1. Bars represent mean ± SEM of two independent replicates. *** p ≤ 0.005 by unpaired Student’s *t-*test. **b)** Linear regression of relative m^6^A enrichment from (a). Points and error bars mark mean ± SEM of two independent replicates. **c)** Change in MeRIP-RT-qPCR vs. MeRIP-seq enrichment for peaks detected as significantly differentially expressed with infection of Huh7 cells by dengue virus, Zika virus, and hepatitis C virus. **d)** Number of replicates of infected vs. uninfected cells needed to detect the peaks in (c). Replicates were randomly subsampled 10 times to calculate the fraction of subsamples in which peaks were called as significant by the GLMs or QNB. Boxes span the 1^st^ to 3^rd^ quartiles, with medians indicated. Whiskers show the minimum and maximum points within ±1.5x the interquartile distance from the boxes. Results for each subsample of replicates are shown as jittered points.

We next assessed the correlation between m^6^A enrichment observed using MeRIP-seq and MeRIP-RT-qPCR using data from our recent work that identified 58 peak changes in m^6^A in Huh7 cells following infection by four different viruses (50). For those experiments, we again selected peaks that change based on results from the union of QNB and the GLM approaches. We found that the magnitude of changes in common among viruses correlated between MeRIP-seq and MeRIP-RT-qPCR, both across peaks (Pearson’s R = 0.57, p = 3.7E-6) and within single peaks across viruses (13 out of 19 peaks showed positive correlations, four of which had p-values < 0.05 with three data points) (**Figure 5c, Additional File 2: Supplementary Figure 5**). Given the correlation we found between MeRIP-seq and MeRIP-RT-qPCR, it is unclear why changes in IP over input sequencing reads were undetectable at the peaks reported by Bertero et al. (2018) and Huang et al. (2019) but differences in peaks were successfully validated using MeRIP-RT-qPCR (22,70). Based on these discrepancies, while MeRIP-RT-qPCR can be used as an initial method of validation for predicted peak changes, additional methods are necessary to confirm quantitative differences in m^6^A levels and to resolve points where the assays do not agree.

We next used our peaks validated with MeRIP-RT-qPCR to estimate the number of replicates necessary for detection of changes with either the GLM or QNB methods. Using a permutation test, we downsampled infected and uninfected replicates and reran statistical detection of changes. We found that approximately 6-9 replicates were necessary for consistent detection (in at least 50% of subsamples) of most peak changes (**Figure 5d**). Schurch et al. (2016) and Conesa et al. (2016) produced similar recommendations for basic RNA-seq studies, finding that 6-12 replicates were necessary to detect most changes in gene expression and that changes of 1.25 were detectable 25% of the time with five replicates, rising to 44% with ten replicates, respectively (36,77). While our findings broadly agree with these recommendations for RNA-seq, they also suggest that almost all published MeRIP-seq studies to date are underpowered.

## Discussion

In the eight years since MeRIP-/m^6^A-seq was first published (16,17), many studies have used these methods to examine the function of m^6^A, its distribution along mRNA transcripts, and how it might be regulated under various conditions. While 35 out of 64 of the MeRIP- and miCLIP-seq papers we surveyed **(Additional File 1: Supplementary Table 1)** refer to m^6^A as “dynamic”, and, by contrast, only two describe the modification as “static”, the literature is unclear on what is meant by the word “dynamic”. There is mixed evidence as to whether m^6^A is reversible through demethylation by the proposed demethylases FTO and ALKBH5 (71,78–80). Recent research using an endoribonuclease-based method for m^6^A detection suggests that ALKBH5 has only a mild suppressive effect on m^6^A levels and FTO no effect (54). Although m^6^A does not appear to change over the course of an mRNA’s lifetime at steady-state (3), whether it changes in response to a particular stimulus and at what point is less clear. Some studies have suggested that m^6^A may be modulated through changes in methyltransferase and demethylase expression, producing consistent directions of change across transcripts (8,23,34), through alternative mechanisms involving microRNA, transcription factors, promoters, or histone marks (21,22,66,70,81), or through indeterminate mechanisms (17,20,25–28,52). However, based on our reanalysis of available MeRIP-seq data, there is still only meagre support for widespread changes in m^6^A across the transcriptome independent of changes in the expression of methylation machinery (e.g. increases or decreases in METTL3 expression).

In particular, replication of peaks and changes in peaks across studies is limited. As with other RNA IP-based methods, MeRIP-seq data contains noise, owing to technical and biological variation (82). In fact, while peak overlaps reach ∼80% between replicates of the same study, they decrease to a median of 45% between studies, most of which use 2-3 replicates each **(Figure 1)**. Given that the detection of peaks is so variable and that peak heights differ among replicates, it is perhaps not surprising that peak changes have yet to be reproduced between multiple studies of similar conditions. Indeed, variability in MeRIP-seq could also mask differences in m^6^A regulation among cell types, which have been described in mouse brains (34) and in cell lines exposed to KSHV (28). To distinguish biological and technical variation, it will therefore be particularly important to test if multiple groups using the same cell line and conditions can better reproduce changes in m^6^A.

Disparities in the methods used to detect changes in m^6^A_(m)_ peaks also play a role in differing conclusions among studies. Here, we analyzed four statistical methods to detect changes in peaks and found that three of these methods showed uniform or conservatively shifted p-value distributions and were able to identify changes in m^6^A_(m)_ independent of changes in gene expression. We therefore suggest that these statistical methods, in combination with filters for input levels in both conditions and the difference in log_2_ fold change between peaks and genes, can be used to identify candidate m^6^A_(m)_ sites from MeRIP-seq data for further analysis and validation (**Figure 6**). Based on our results, while MeTDiff works for peak detection, we do not recommend MeTDiff for peak change detection as it does not control well for differences in gene expression **(Figure 3)**. Similar to others (33), we found that plotting predicted m^6^A changes was invaluable and that appropriate scaling for gene coverage could reveal changes proportional to gene expression. In addition, plotting the standard deviation in transcript coverage can help assess typical variation in peak height among replicates. We note that both differential methylation of a gene and methylation of a gene that is differentially expressed could be important, but they should not be conflated when considering the role of m^6^A in transcript regulation.

**Figure 6:**
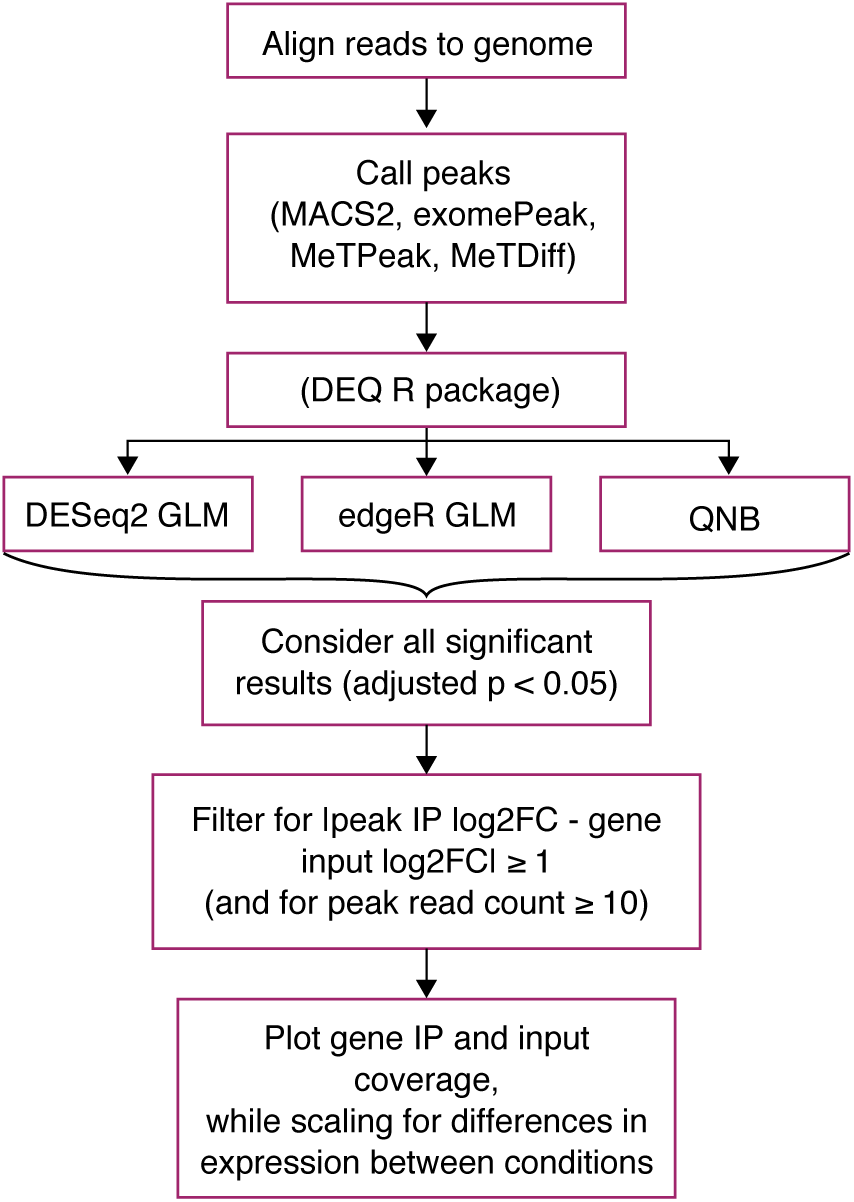
Proposed approach to identify candidates for m^6^A_(m)_ changes for further validation using MeRIP-seq data. We suggest predicting changes in m^6^A_(m)_ using DESeq2, edgeR, and QNB, and have implemented the DEQ package in R to facilitate this.

The extent to which m^6^A changes on particular transcripts and whether it changes in binary presence/absence or in degree is unclear. MeRIP-RT-qPCR could detect methylation differences in in vitro transcribed RNA. Further, we found that these changes correlated with differences in MeRIP-seq enrichment. However, neither MeRIP-seq nor MeRIP-RT-qPCR can reveal the precise fraction of transcript copies modified by m^6^A. In general, antibody-based methods are subject to biases, including from differences in binding efficiencies based on RNA structure and motif preferences (83). There is an oft-cited but little-used method for quantification of m^6^A, site-specific cleavage and radioactive-labeling followed by ligation-assisted extraction and thin-layer chromatography (SCARLET) (19). However, this method can be challenging, works only for highly abundant transcripts, and is impractical for transcriptome-wide analysis. A recently developed endoribonuclease-based, antibody-independent approach for m^6^A detection is promising in terms of quantification of m^6^A, but its use is limited to a subset of m^6^A sites within DRAC motifs ending in ACA (∼16% of all sites) (44,45). So far, comparison to this data suggests that antibody-based approaches may underestimate the number of m^6^A sites (54). Alternative methods to detect m^6^A based using single-molecule sequencing (including direct RNA sequencing and real-time cDNA synthesis) are under development and may offer ways to detect, quantify, and phase m^6^A sites, but these have not yet been shown to accurately detect m^6^A across a cellular transcriptome (84–86). For now, site-specific SCARLET is the only option to biochemically validate proposed changes in m^6^A at most motifs.

## Conclusions

Our work reveals the limits of MeRIP-seq reproducibility for the detection of m^6^A_(m)_ and in particular suggests caution when using MeRIP-seq for the detection of changes in m^6^A_(m)_. To increase confidence in predicted changes in m^6^A_(m)_, we propose statistical approaches that account for differences in gene expression between conditions and variability among replicates. These methods can be used to gain insight into the regulation and function of m^6^A_(m)_ and to predict specific sites for validation before the development of high-throughput alternatives to MeRIP-seq, and similar strategies may be applicable to other types of RNA sequencing assay.

## Methods

### New MeRIP-seq data

#### Huh7 data

Total RNA was extracted from Huh7 cells using Trizol (Thermo-Fisher). mRNA was purified from 200 μg total RNA using the Dynabeads mRNA purification kit (Thermo-Fisher) and concentrated by ethanol precipitation. Purified mRNA was fragmented using the RNA Fragmentation Reagent (Thermo-Fisher) for 15 minutes followed by ethanol precipitation. Then, MeRIP was performed using EpiMark N6-methyladenosine Enrichment kit (NEB). 25 μL Protein G Dynabeads (Thermo-Fisher) per sample were washed three times in MeRIP buffer (150 mM NaCl, 10 mM Tris-HCl, pH 7.5, 0.1% NP-40) and incubated with 1 μL anti-m^6^A antibody (NEB) for 2 hours at 4°C with rotation. After washing three times, anti-m^6^A conjugated beads were incubated with purified mRNA with rotation at 4°C overnight in 300 μL MeRIP buffer with 1 μL RNAse inhibitor (recombinant RNasein; Promega). Beads were then washed twice with 500 μL MeRIP buffer, twice with low salt wash buffer (50 mM NaCl, 10 mM Tris-HCl, pH 7.5, 0.1% NP-40), twice with high salt wash buffer (500 mM NaCl, 10 mM Tris-HCl, pH 7.5, 0.1% NP-40), and once again with MeRIP buffer. m^6^A-modified RNA was eluted twice in 100 μL MeRIP buffer containing 5mM m^6^A salt (Santa Cruz Biotechnology) for 30 minutes at 4°C with rotation and concentrated by ethanol precipitation. RNA-seq libraries were prepared from eluate and the 10% of RNA set aside as input using the TruSeq mRNA library prep kit (Illumina) and checked for fragment length using the Agilent 2100 Bioanalyzer. Single-end 50 base pair reads were sequenced on an Illumina HiSeq 2500.

#### Heat shock

Early passage OCI-Ly1 diffuse large B-cell lymphoma cells were grown in Iscove’s modified Eagle Medium (IMDM) with 10% fetal bovine serum (FBS). OCI-Ly1 cells were obtained from the Ontario Cancer Institute and regularly tested for *Mycoplasma* contamination by PCR and identified by single nucleotide polymorphism. Cells were maintained with 1% penicillin/streptomycin in a 37°C, 5% CO_2_, humidified incubator. In these growing conditions, heat shocked cells were exposed to 43 °C for 1 hour, followed by 1 hour of recovery at 37°C while control cells were maintained at 37°C. Following treatment, cells were processed at 4°C to obtain total cell lysates. Lysates were immunoprecipitated for m^6^A_(m)_ using Synaptic Systems antibody (SYSY 202 003) following the protocol described in Meyer, et al (2012) and sequenced on an Illumina HiSeq 2500 (16).

### Read processing

Reads were trimmed using Trimmomatic (87) and aligned to the human genome (hg38) or the mouse genome (mm10), as appropriate, using STAR, a splice-aware aligner for RNA-seq data (72). We used the flag “--outFilterMultimapNmax 1” to keep only uniquely aligned reads. Scripts used for alignment are provided with the rest of the analysis scripts at https://github.com/al-mcintyre/merip_reanalysis_scripts.

### Peak detection and comparison

IP over input peaks were called using MACS2 callpeak using the parameters “--nomodel --extsize 100 (or, if available, the approximate fragment size for a specific experiment to extend reads at their 3’ ends to a fixed length) --gsize 100e6 (the approximate size of mouse and human transcriptomes based on gencode annotations)” (49). No filter for coverage was applied at the stage of peak detection. Transcript coverage was estimated using Kallisto (88) with an index construct 31mers, except for the Schwartz et al (2014) data set, where the reads were too short and an alternative index based on 29mers was constructed (33). For **Figure 1b**, the full union of unique peaks was taken and the percent of that set detected in single replicates calculated. Intersects between peaks that overlapped for transcripts with ≥10X mean coverage in both samples were taken using bedtools (89) for **Figure 2**, allowing a generous minimum of 1 overlapping base. Heatmaps for peak overlaps were generated using the ComplexHeatmap package in R (90). MeRIP-seq data sets in **Figure 2b** included those for human cell lines in **Figure 2a**, other data sets from the same studies and any data sets that shared the same cell lines, and other data sets that looked at multiple human cell types. We considered only data sets from baseline conditions in **Figure 2** (untreated cells and knockdown controls).

### Poisson and negative binomial fits

Reads aligned to peaks were counted using featureCounts from the Rsubread package (91). Poisson and negative binomial models were fit to input and IP read counts at peaks using maximum likelihood estimation. Simulated read counts were generated with Poisson or negative binomial distributions based on estimated parameters from the sample, with 500 random generations per model. The log likelihood of seeing read counts from the sample and the simulations given the model parameters was then calculated and the mean taken across all peaks.

### Peak change detection and generalized linear models

Generalized linear models to detect changes in IP coverage while controlling for differences in input coverage were implemented based on a method previously applied to HITS-CLIP data (58). Full and reduced models were constructed as follows:

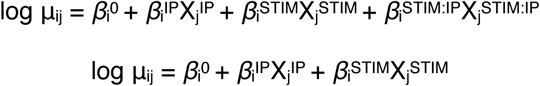

Where μ_ij_ is the expected read count for peak i in sample j, modelled as a negative binomial distribution, X_j_^IP^ = 1 for IP samples and 0 for input samples, and X_j_^STIM^ = 1 for samples under the experimental intervention and 0 for control samples.

Statistical significance was then assessed using a chi-squared test (df=1) for the difference in deviances between the full and reduced models, with the null hypothesis that the interaction term (β_i_STIM:IP) for differential antibody enrichment driven by the experimental intervention is zero. The likelihood ratio test was implemented through DESeq2 (56) and edgeR (57), two programs developed for RNA-seq analysis that differ in how they filter data and in how they estimate dispersions for negative binomial distributions. Generalized linear models implemented through edgeR included a term for the normalized library size of sample j.

QNB was run as suggested for experiments with biological replicates, where each IP and input variable (“ip1”, etc.) consisted of a matrix of peak counts for either condition 1 or condition 2: > qnbtest(ip1, ip2, input1, input2, mode=“per-condition”)

We extracted functions from MeTDiff so that we could supply our own peaks and thus control for differences in peak detection among tools. The main post-peak calling function, diff.call.module, was run as follows using the same count matrices as for QNB: > diff.call.module(ip1, input1, ip2, input2)

Gene and peak expression changes were estimated as log_2_ fold changes from DESeq2 based on differences in input read counts aligned to genes and IP read counts aligned to peaks, respectively, and the change in peak relative to gene enrichment was calculated as the absolute difference in log_2_ fold change between those values.

### Comparison to published studies

The sources for published estimates of m^6^A peak changes included in our comparison are listed in **Additional File 1: Supplementary Table 4**. Significant (FDR-adjusted p < 0.05) peaks were considered for DESeq2, edgeR, and QNB, run as described above. We also considered a filtered set of peaks derived from the union of significant peaks from the three tools with additional filters for location within exons, |log_2_ fold change between peak IP and gene input| ≥ 1, and a minimum peak read count of 10 across replicates and conditions. We used gProfiler to calculate enrichment of functional categories (92).

In **Figure 4b-c**, we selected *Hspa1a/HSPA1A* as our representative gene for heat shock because it was the primary example cited by Zhou et al. (2015) and Meyer et al. (2015) (4,20). For HIV, we selected *PSIP1* because it was among the 56 genes reported by Lichinchi et al. (2016a) (25), it plays a known role in HIV infection, and we detected a peak in the gene using MACS2.

For KSHV, we compared significant results (adjusted p < 0.05) from QNB and GLMs (DESeq2 and edgeR), with additional filtering for |peak IP – gene input log_2_ fold change| ≥ 1 (lowering this threshold to 0.5 did not change results), for data from Hesser et al. (2018) (27) in lytic vs. latent iSLK.219 cells and data from Tan et al. (2018) (28) in lytic vs. latent iSLK BAC16 cells. We used the same approach to compare data from Rubio et al. (2018) and Winkler et al. (2019) (75,76) for response to dsDNA. Data sets used for site-specific comparisons are summarized in **Additional File 1: Supplementary Table 5**.

Gene coverage was plotted using CovFuzze (https://github.com/al-mcintyre/CovFuzze), which summarizes mean and standard deviation in coverage across available replicates (93). Pearson’s correlations were taken for **Supplementary Figure 4** for peaks expressed above a minimum input peak read count of 10 across replicates and conditions.

### Spike-in controls and MeRIP-RT-qPCR

In vitro transcribed (IVT) controls were provided by the Jaffrey Lab and consisted of 1001 base long RNA sequences with three adenines in GAC motifs (**Additional File 1: Supplementary Table 6**) either fully methylated or unmethylated. m^6^A and A controls were mixed in various ratios (1:9, 3:7, and 9:1) that approximate the variation in m^6^A levels detected by SCARLET (m^6^A levels at specific sites have been reported to vary from 6-80% of transcripts (19)). Modified and unmodified standards were mixed at the indicated ratios to yield a final quantity of 0.1 fmol, 1 fmol, and 10 fmol. Mixed RNA standards were added to 30 μg total RNA from Huh7 cells, along with 0.1 fmol of positive (m^6^A-modified *Gaussia* luciferase RNA, “GLuc”) and negative control (unmodified *Cypridina* luciferase, “CLuc”) spike-in RNA provided with the N6-methyladenosine Enrichment kit (EpiMark). Following MeRIP as described above, cDNA was synthesized from eluate and input samples using the iScript cDNA synthesis kit (Bio-Rad), and RT-qPCR was performed on a QuantStudio Flex 6 instrument. Data was analyzed as a percent of input of the spike-in RNA in each condition relative to that of the provided positive control spike-in.

For MeRIP-RT-qPCR to test peak callers, Huh7 cells plated in 6-well plates were transfected with siRNAs against METTL3 and METTL14 (Qiagen; SI04317096 and SI00459942) or non-targeting control siRNa (SI03650318) using Lipofectamine RNAiMax (Thermo Fisher) twice, 24 hours apart. 48 hours following the second round of siRNA transfection, cells were harvested in TRIzol reagent and total RNA was extracted. 30 μg total RNA was fragmented for 3 mins at 75°C, concentrated by ethanol precipitation, and MeRIP-RT-qPCR was performed as described above. Primers used for RT-qPCR are provided in **Additional File 1: Supplementary Table 7** and siRNA sequences in **Additional File 1: Supplementary Table 8.**

### Cell culture and infection (data used for MeRIP-RT-qPCR comparisons)

Huh7 cells were grown in DMEM (Mediatech) supplemented with 10% fetal bovine serum (HyClone), 2.5 mM HEPES, and 1X non-essential amino acids (Thermo-Fisher). The identity of the Huh7 cell lines was verified using the Promega GenePrint STR kit (DNA Analysis Facility, Duke University), and cells were verified as mycoplasma free by the LookOut Mycoplasma PCR detection kit (Sigma). Infectious stocks of a cell culture-adapted strain of genotype 2A JFH1 HCV were generated and titered on Huh7.5 cells by focus-forming assay (FFA), as described (94). Dengue virus (DENV2-NGC), West Nile virus (WNV-NY2000), and Zika virus (ZIKV-PRVABC59) viral stocks were generated in C6/36 cells and titered on Vero cells as described (94). All viral infections were performed at a multiplicity of infection of 1 for 48 hours.

## Supporting information

Additional file 3: Results with MeTDiff peak caller.

Additional file 4: Peaks and statistical test results for experiments in Figures 3-4.

Additional file 5: Response to reviewers.

## Declarations

### Ethics approval and consent to participate

Not applicable

### Consent for publication

Not applicable

## Availability of Data and Materials

MeRIP-seq data for the Huh7 negative controls is available in the GEO repository, under accession number GSE130891. MeRIP-seq data for heat shock in B-cell lymphoma is available under accession number GSE130892. Accession numbers for all other data sets reanalyzed in the study are included in Additional File 1: Supplementary Tables 1-5. Scripts used for analysis are available at https://github.com/al-mcintyre/merip_reanalysis_scripts and a pipeline implementing generalized linear models through DESeq2 and edgeR, as well as QNB, is provided at https://github.com/al-mcintyre/deq.

## Authors’ contributions

A.B.R.M. and C.E.M. conceived the study. A.B.R.M. developed and ran the analyses and wrote the manuscript with N.S.G. and S.M.H. S.R.J. provided in vitro controls for MeRIP-RT-qPCR. N.S.G. prepared MeRIP-seq libraries for the Huh7 controls and ran MeRIP-RT-qPCR tests. L.C. contributed additional heat shock data. All authors read and edited the manuscript.

## Acknowledgements

We would like to thank Christina Leslie and Jonathan Victor for statistical advice, Aashiq Mirza for the MeRIP-RT-qPCR controls, and Helen Lazear for WNV infection. We would also like to thank the Epigenomics Core at Weill Cornell for preparing MeRIP-seq libraries, the Scientific Computing Unit (SCU), and New England Biolabs for donating anti-m^6^A antibodies.

---

**Additional file 1:** Supplementary Tables. (xls)

**Additional file 2:** Supplementary Figures. (pdf)

**Additional file 3**: Results with MeTDiff peak caller. (pdf)

**Additional file 4**: Peaks and statistical test results for experiments in Figures 3-4. (zip)

**Additional file 5**: Response to reviewers. (txt)

