## Additional file 2: Supplementary Figures. for "Limits in the detection of m^6^A changes using MeRIP/m^6^A-seq"

### Supplementary Figure 1

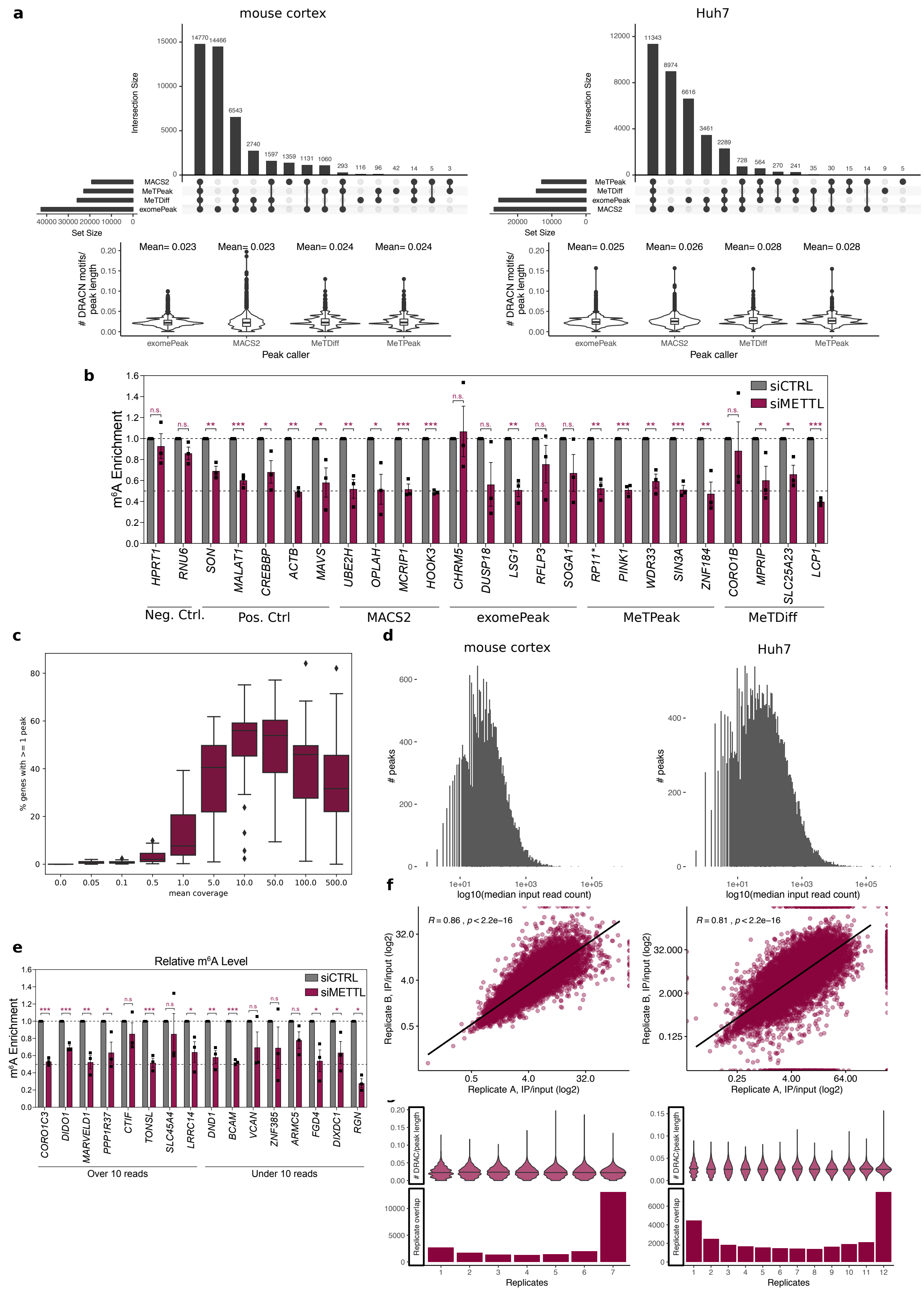

**Supplementary Figure 1**: **a)** Peak overlaps across peak calling methods (top) and enrichment of DRAC motifs (the number of DRAC motifs divided by peak length, with mean indicated above the violin plots) across methods. Boxes within the violin plots span the $1^{st}$ to $3^{rd}$ quartiles, with medians indicated. Whiskers show the minimum and maximum points within $\pm$1.5x the interquartile distance from the boxes, with points beyond that range shown as outliers. **b)** MeRIP-RT-qPCR analysis of a subset of peaks called by single tools to test for METTL3-METTL14 dependence. Huh7 cells were treated with siRNAs targeting *METTL3* and *METTL14* (siMETTL) or non-targeting control siRNA (siCTRL) and MeRIP-RT-qPCR was performed using primers under uniquely called peaks. HPRT1 (known unmethylated transcript) and RNU6 (known to be methylated by METTL16) were negative controls. Positive controls were previously published known methylated transcripts. **c)** The percent of genes with ≥ 1 m6A peak for various mean coverage bins (0-0.05, 0.05-0.1, etc.). Results are summarized across the experiments included in Figures 1 and 2. Boxplot center lines show medians and whiskers 1.5x interquartile range. **d)** Histograms of peaks detected at increasing median input levels across replicates confirms that peak detection requires adequate input coverage (here, few peaks are called at input levels below ten). **e)** METTL3/14-dependence of a random selection of peaks detected in low expression < 10 reads vs. higher expression > 10 reads sites from Huh7 data. MeRIP-RT-qPCR analysis of relative m$^6$A level RNA harvested from Huh7 cells that were treated with siCTRL or siMETTL3/14. Values are the mean ± SD of 3 replicates. * $p < 0.05$, ** $p < 0.01$, *** $p < 0.001$ by unpaired Student's t test. n.s = not significant. **f)** The correlation between log$_2$ fold IP/input read counts between two random replicates. **g)** The enrichment of DRAC motifs calculated as # DRAC motifs/peak length for peaks detected by X replicates shows little change for peaks detected in multiple replicates, with the number of peaks detected by X replicates shown below. The lines in the violin plots indicate the median. **a,d,f,g** show data from mouse cortex (left) and Huh7 cells (right) at baseline conditions.

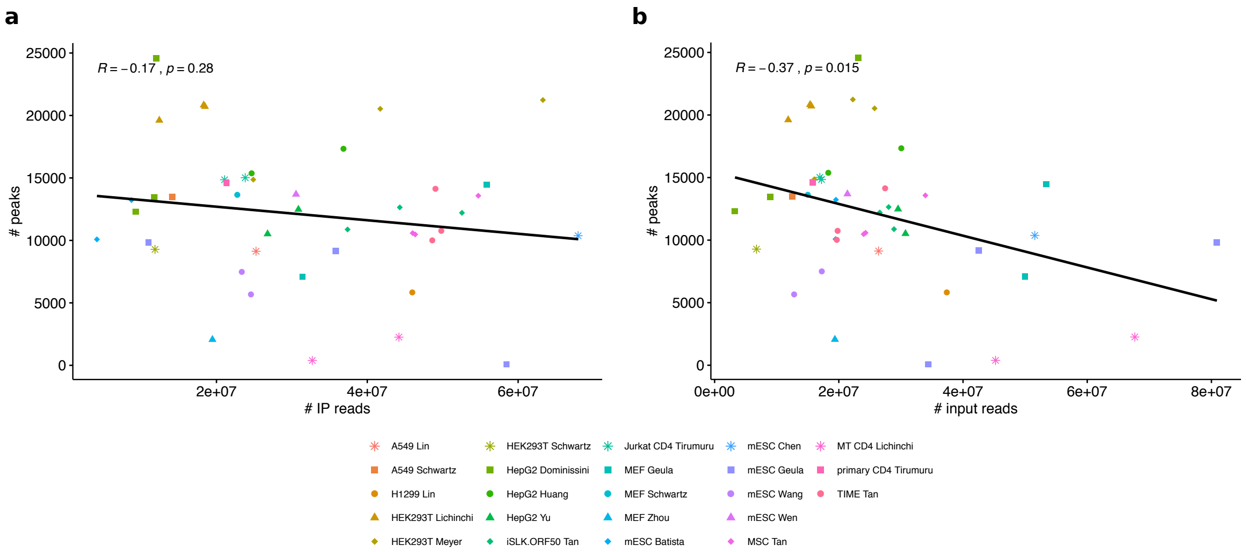


**Supplementary Figure 2. a-b)** The number of peaks detected per replicate as a function of the number of IP (a) and input (b) sequencing reads for the studies included in **Figure 2** (**Supplementary Table 2**).

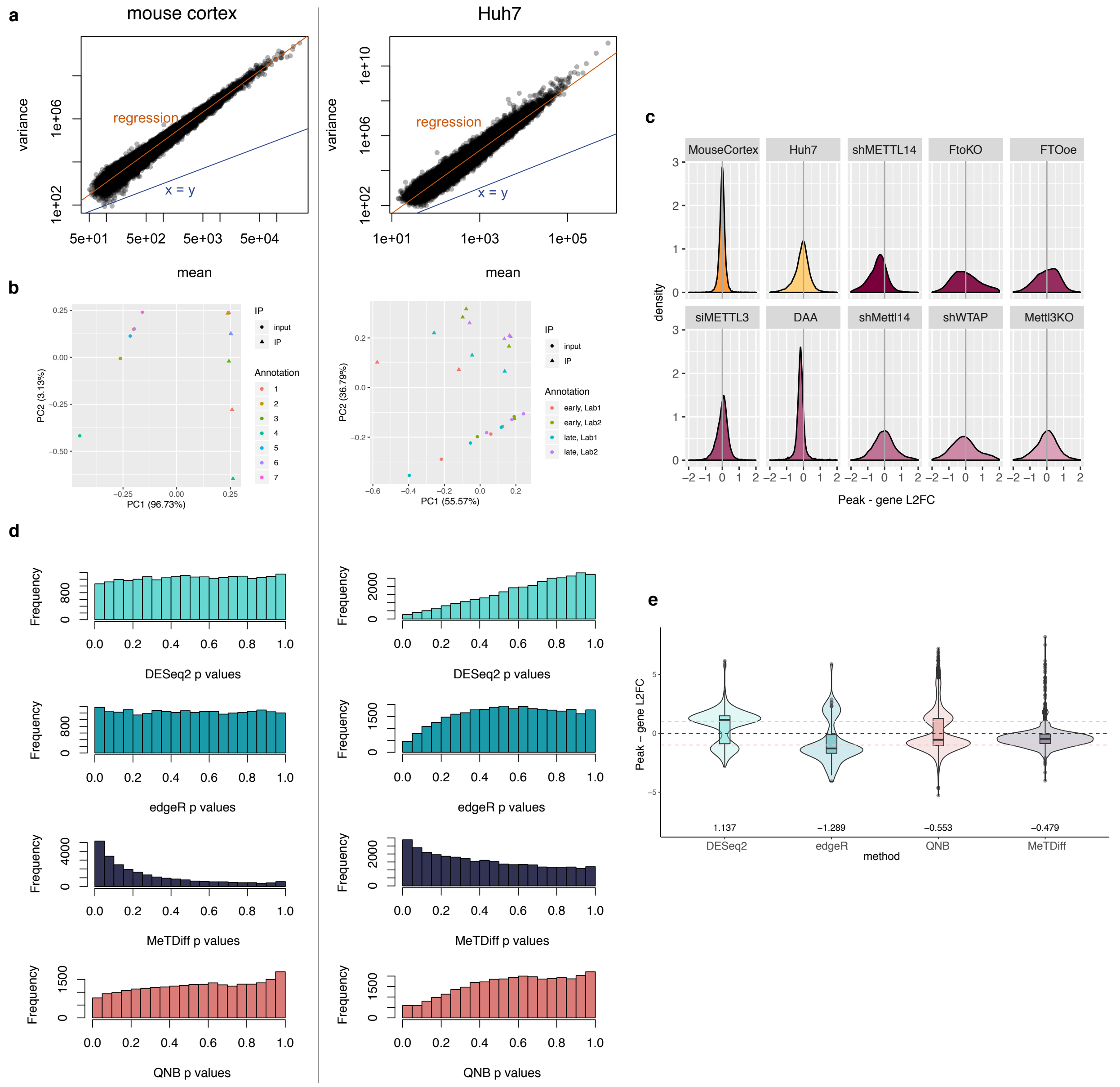


**Supplementary Figure 3**: **a)** The variance of input and IP read counts under peaks exceeds the mean (mean-variance equivalence line shown in blue, linear regression fit shown in orange), indicating overdispersion. **b)** The top two principal components for IP and input read counts for the two negative control experiments. **c)** The distribution of peak – gene $\log_2$ fold changes for the positive and negative control experiments across all genes. **d)** P-value distributions indicate conservative shifts for the generalized linear models and QNB for the Huh7 data set (right) but are uniform as expected for the mouse cortex data set (left). MeTDiff p-value distributions show an excess of significant peaks in the negative controls. **e)** The distributions of peak-gene $\log_2$ fold changes for significant peaks called in the positive control experiments shows depletion where peak and gene $\log_2$ fold changes are equivalent for all methods except MeTDiff. Dashed lines indicate differences in $\log_2$ fold change of -1, 0, and 1 and medians are summarized below the distributions. Boxes within the violin plots span the $1^{st}$ to $3^{rd}$ quartiles, with medians indicated. Whiskers show the minimum and maximum points within $\pm 1.5$ times the interquartile distance from the boxes, with points beyond that range shown as outliers.

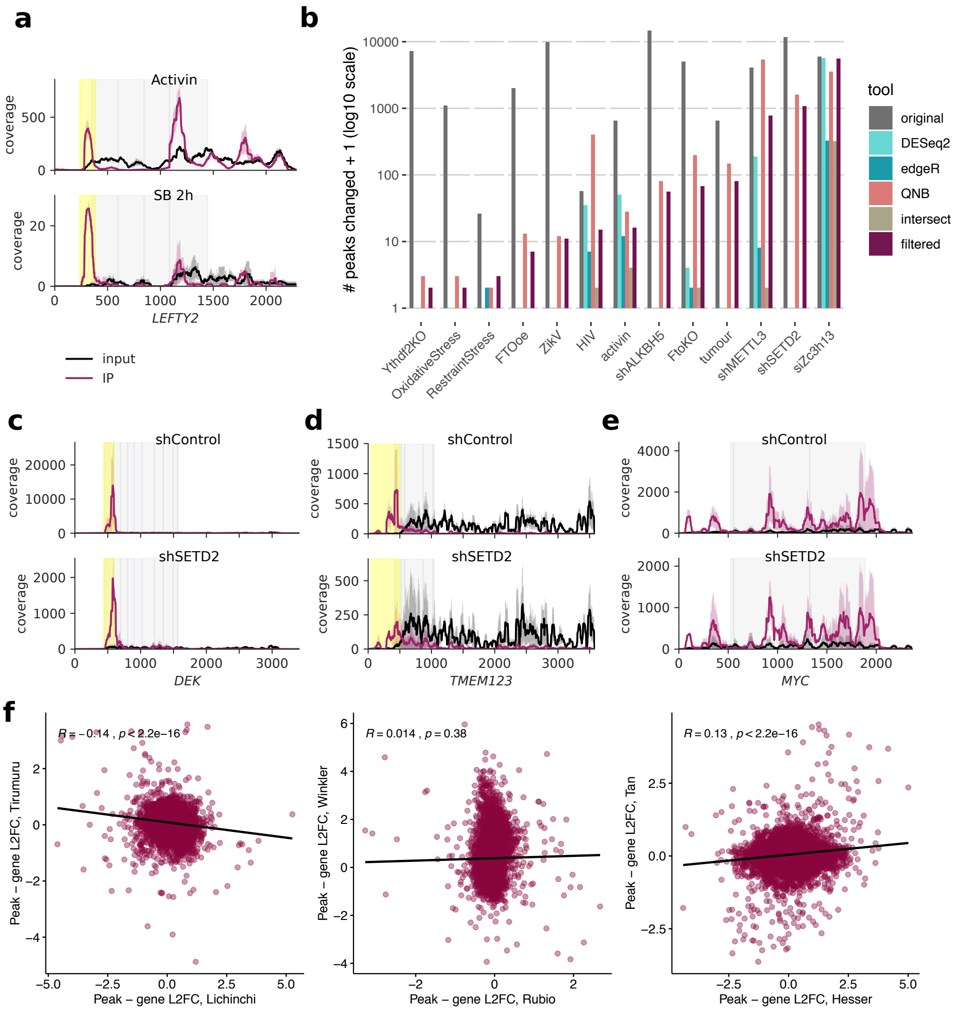


**Supplementary Figure 4: a)** Coverage for a gene involved in pluripotency that we detected as significantly differentially methylated with activin-NODAL inhibition by SB431542 (SB). The peak did not pass the filter for minimum input reads. **b)** Detected $m^6A_{(m)}$ changes in thirteen published data sets that measured $m^6A_{(m)}$ peak changes between two conditions **(Additional File 1: Supplementary Table 4)**. The number of peaks detected as changed in the original published analyses are compared to the number of peaks with FDR-adjusted p-values < 0.05 in our reanalysis using DESeq2, edgeR, or QNB, and taking the union of results from these three tools with additional filters for $\log_2$ fold difference in peak and gene changes of ≥1 ("filtered"), without an additional filter for input read counts. **c)** The sole peak detected as changed in the Huang et al. (2019) shSETD2 data set that passed a minimum read threshold. **d)** The most significant peak predicted by QNB in the Huang et al. (2019) shSETD2 data set. **e)** A gene reported as hypomethylated by Huang et al. (2019) based on MeRIP-seq and MeRIP-RT-qPCR data in which we did not detect any significant changes in methylation, with or without a filter for input reads. In general, low input coverage in the first two replicates from this study appeared to contribute to the elevated standard deviations in coverage. **f)** The correlation between changes in enrichment (Δ peak - gene $\log_2$ fold change between conditions) at the same sites between experiments that studied similar conditions.

**a,c-e)** Lines show the mean coverage across three replicates for input and IP samples, while shading shows the standard deviation. Peaks detected as significantly changed are highlighted in yellow and coding sequences are shown in grey.

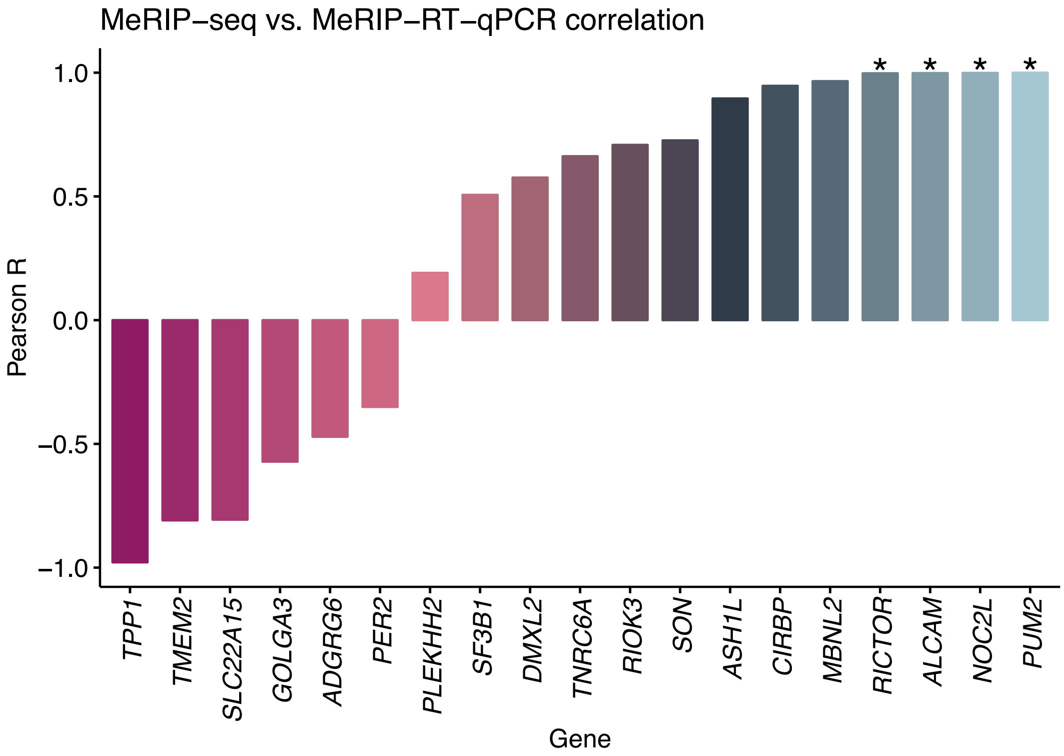


**Supplementary Figure 5:** Pearson correlation between the change in MeRIP-RT-qPCR and change in MeRIP-seq enrichment for specific peaks (labelled by gene name) detected as significantly differentially expressed with infection of Huh7 cells by dengue virus, Zika virus, and hepatitis C virus. Asterisks indicate correlations with $p < 0.05$.
