## Additional file 3: Results with MeTDiff peak caller. for "Limits in the detection of m^6^A changes using MeRIP/m^6^A-seq"

### Additional File 3: Results using MeTDiff peak caller

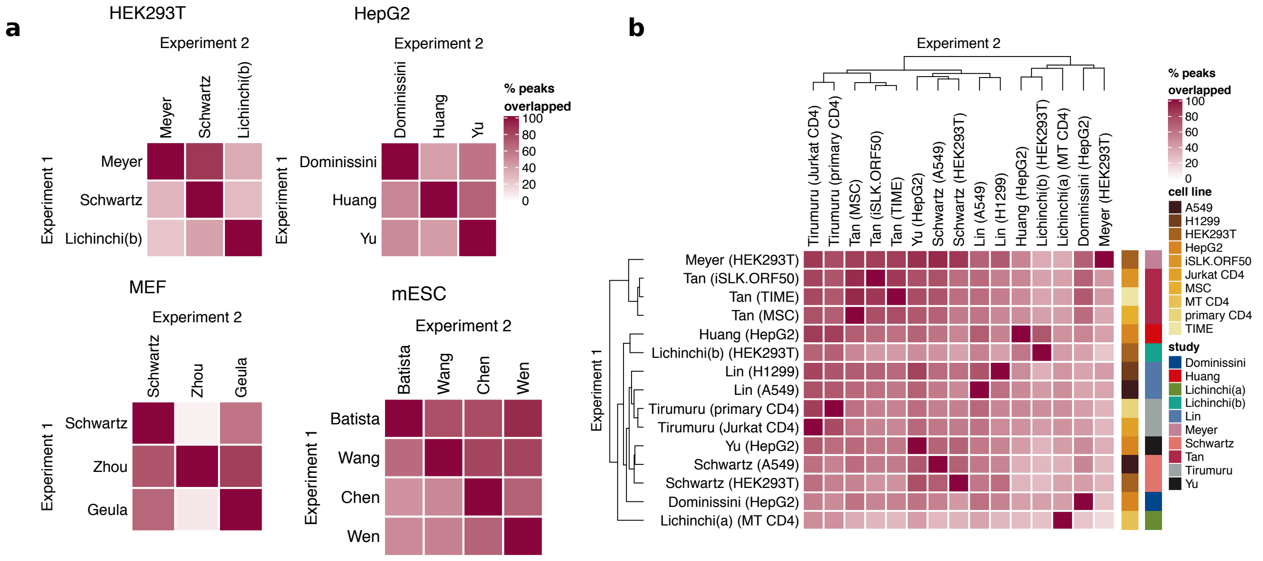


**Supplementary Figure 1**: **a)** Peak detection using MeTDiff between studies that used the same cell type shows variable overlap. Overlap was calculated as the percent of peaks detected in Experiment 1 with an overlap of ≥ 1 base pair with peaks from Experiment 2 (compare to Figure 2a), **b)** Peak detection across tissue and cell types shows samples from the same study cluster better together than samples from the same tissue (compare to Figure 2b). Median overlap was 55%.

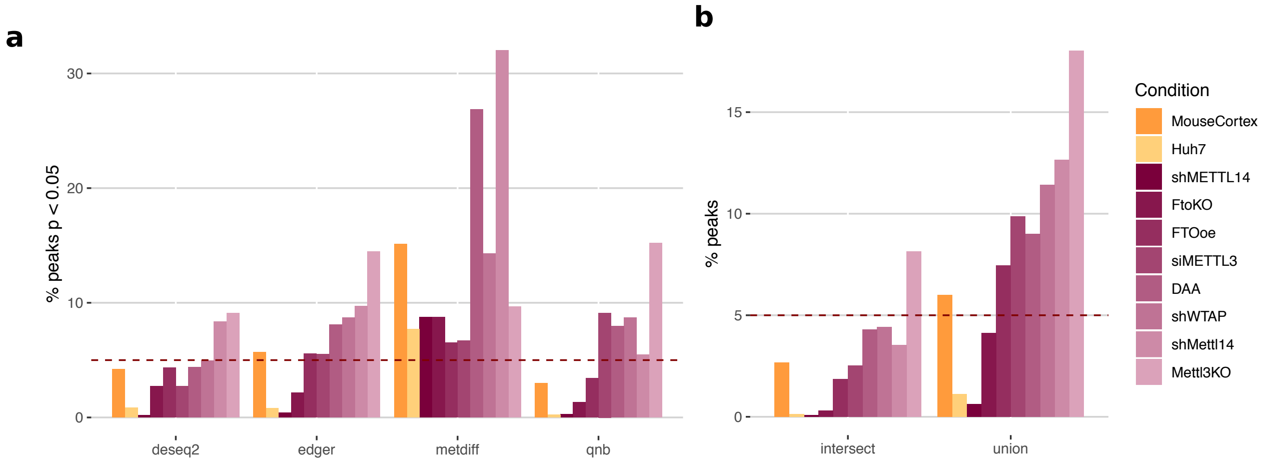


**Supplementary Figure 2**: **a)** The percent of sites below an unadjusted p-value threshold of 0.05 for different methods (described in **Table 1**) to detect differential methylation in negative controls between two groups at baseline conditions and positive controls in which methylation processes were disrupted with respect to baseline conditions (**Additional File 1: Supplementary Table 3**). The line at 5% indicates the expected proportion of sites given a uniform p-value distribution (see **Additional File 2: Supplementary Figure 3c**), while colours indicate negative (orange) and positive (purple) control experiments (compare to Figure 2b). **b)** The intersect and union of peaks with $p < 0.05$ from DESeq2, edgeR, and QNB from (a), coloured as in (a).

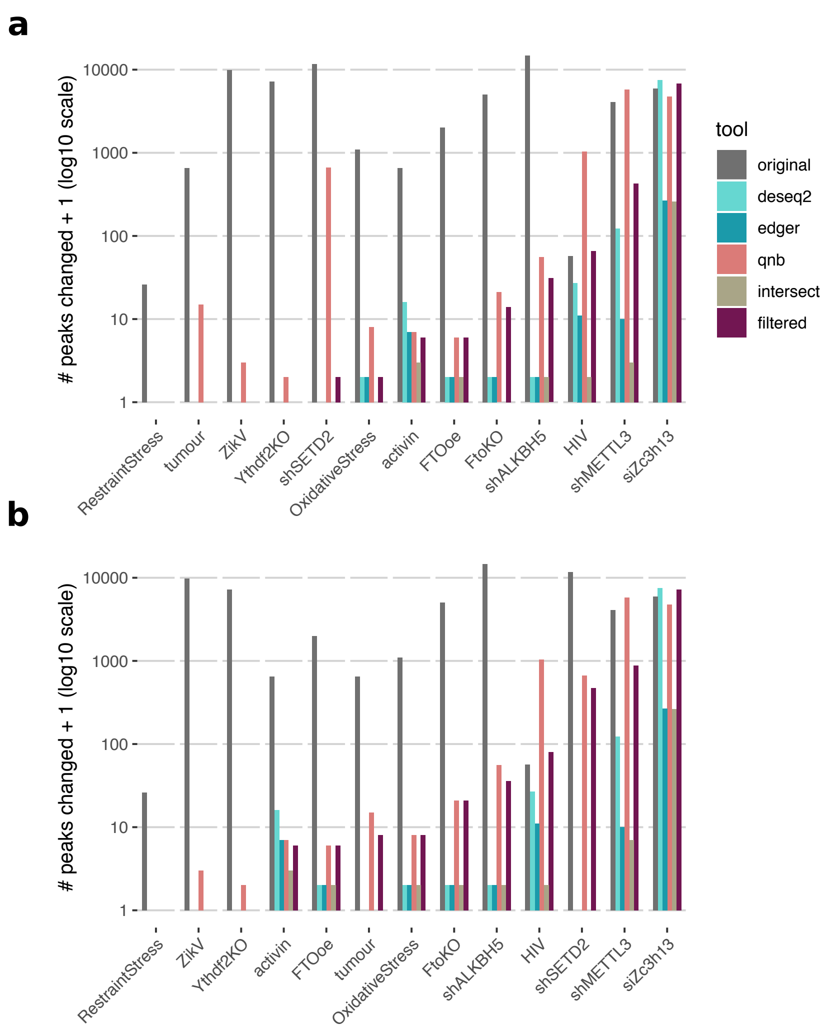


**Supplementary Figure 3: a)** Detected m$^6$A$_{(m)}$ changes in thirteen published data sets that measured m$^6$A$_{(m)}$ peak changes between two conditions **(Additional File 1: Supplementary Table 4)**. The number of peaks detected as changed in the original published analyses are compared to the number of peaks with FDR-adjusted p-values < 0.05 in our reanalysis using DESeq2, edgeR, or QNB, and taking the union of results from these three tools with additional filters for log$_2$ fold difference in peak and gene changes of ≥1 and peak read counts ≥10 across all replicates and conditions ("filtered") (compare to **Figure 4a**), **b)** The same, without a threshold for peak read counts (compare to **Additional File 2: Supplementary Figure 4b**).
