## Additional file 5: Response to reviewers. for "Limits in the detection of m^6^A changes using MeRIP/m^6^A-seq"

### Reviews and Responses

*The paper was rejected based on these reviews. We are happy to post them along with our full responses (in blue) in case others have similar questions (please also feel free to comment on bioRxiv #openpeerreview). Needless to say, we disagree with these evaluations and aim to provide useful insight into the issues with MeRIP-seq data and conclusions that rely on it, rather than the ideal pipeline or analysis tool. Good analysis can't fix bad data, so validation is always key. -AM*

**Reviewer #1:**

The manuscript by McIntyre et al. tried to estimate reproducibility across MeRIP/m6A-seq experiments from different publications using a computational workflow defined by the authors. However, the authors chose a ChIP-seq data/genome-based software (MACS2) to call peaks from RIP-seq/transcriptome. Given the different statistical analysis models between ChIP-seq/genome and RIP-seq/transcriptome for calling the peaks, it is of particular concern that the authors rush to some erroneous conclusions which may mislead biologists to use inappropriate software to call peaks from MeRIP/m6A-seq data (details see major comment 1). Moreover, the manuscript used arbitrary parameters to filter peaks called by different software (details see major comment 2). Furthermore, three softwares have been suggested by the authors to analyze differential peaks, but yielded different results. Importantly, although the manuscript focuses on re-analyzing the published MeRIP-seq data, the detailed information about the methods and parameters employed in each manuscript remains totally unclear. Overall, this study has many defects (listed below) and would not be appropriate for publication in this journal.

We thank the reviewer for the comments and careful review, and we think many of these described limitations are readily addressed by this revision and our updated review of the cited work (e.g. most papers used MACS/MACS2, and we also now test a second method). Also, we clarify some information that may have not been as obvious in the first submission. Please see below.

Major comments:

1. The authors claimed that they used MACS2 for peak calling. But as far as I know, MACS2 was designed for genomic sequencing data like ChIP-seq without taking into consideration of the RNA features, such as introns and exons. Moreover, the statistical analysis models are different between ChIP-seq and MeRIP/m6A-seq data. In addition, the MACS slides the window in genome but not in transcript/gene, so the peaks identified by MACS2 may be far beyond the transcripts and are not suitable for downstream analysis. In fact, in Supplementary Figure 1a, MACS2 had the minimal and maximum number of peaks in mouse cortex and Huh7 data, respectively, which further means that MACS2 is not a robust and suitable approach for m6A peak calling.

We thank the reviewer for this comment, which inspired some further experiments and analyses. Based on these results, as well as our summary of the literature (**Supplementary Table 1**), we find that MACS2 is a suitable peak caller and that using an alternative peak caller does not affect our conclusions. We outline our reasons here:

First, the reviewer is correct that MACS2 was designed for genomic sequencing data (and we note this in the text), but it can also be used for analyzing RNA data, as others have previously shown (e.g. Antanaviciute et al., 2017 (1)). In fact, 20 of the MeRIP-seq studies we surveyed in **Supplementary Table 1** used MACS2 or its precursor MACS, several after running their own comparisons to other peak callers (e.g. Engel et al., 2018 (2)). In these studies, papers often filter for exonic peaks after peak calling to establish a set of peaks appropriate for downstream analysis, however, we note that this could also remove true $m^6A$ sites in introns and un-annotated transcripts (according to Ke et al., 2017, these are rare (3)). Our own results in **Supplementary Figure 1a** suggest that the peak callers we tested achieve comparable accuracy in the prediction of $m^6A$ sites (based on equal enrichment of the canonical motif DRAC).

Despite the prevalent use of MACS2 for MeRIP-seq analysis, we still appreciate the reviewer's concern over its use and the need for accurate peak calling. Therefore, we experimentally tested $m^6A$ peaks predicted by the peak callers compared in **Supplementary Figure 1** (MACS2, MeTPeak, MeTDiff, and exomePeak). Specifically, we selected a set of random peaks among those detected by single peak callers in our Huh7 cells for MeRIP-RT-qPCR validation of METTL3/14-dependence. We found that 4/4 MACS2 peaks, 5/5 MeTPeak peaks, and 3/4 MeTDiff peaks showed less enrichment with knockdown of METTL3/14, suggesting these are METTL3/14-dependent $m^6A$ sites (new **Supplementary Figure 1b**). By contrast, we were able to validate only 1/5 of the peaks uniquely identified by exomePeak, although this may have been due to outliers or other experimental variables. These new data are limited but support the use of MACS2 as a peak caller for MeRIP-seq data.

Taking into account our new data and the results from Antanaviciute et al. (2017) discussed above, we have amended the text lines 121-129 to read: "we assessed the METTL3/METTL14-dependence of specific peaks identified by single tools using MeRIP-RT-qPCR. We found that of these peaks, 4/4 from MACS2, 5/5 from MeTPeak, and 4/5 from MeTDiff showed decreased $m^6A_{(m)}$ enrichment following METTL3/METTL14 depletion, suggesting that these are true $m^6A$ sites. By comparison, only 1/5 of the peaks uniquely called by exomePeak showed statistically significant decreases ($p < 0.05$), although replicate variance was high and 4/5 showed a downward trend (Additional File 2: Supplementary Figure 1b). Since MACS2 is the most commonly used tool for peak calling and was previously found to perform well in comparison with a graphical user interface tool and several other peak callers (51), we used MACS2 for the remainder of our analyses."

Finally, to further validate our results, we re-ran all of the main analyses shown in Figures 2, 3, and 4 using the MeTDiff peak caller instead of MACS2 and summarize the new results in **Additional File 3**. As further support for the broad biological and technical conclusions of our study, we found that none of our primary conclusions changed. Specifically, when using MeTDiff as the peak caller, we also found that: (1) peaks for MeRIP-seq experiments clustered more by study than by cell type or tissue, (2) peak change detection (the step after peak detection to compare enrichment between two conditions) using MeTDiff again appears to detect more false positives than other methods, and (3) most MeRIP-seq studies of various conditions have still likely over-reported peak changes based on comparisons to our statistical approaches.

2. The authors used a coverage threshold of input reads (read counts ≥ 10) to detect m6A peaks, which appears to be the lack of statistical significance and too arbitrary. The sequencing depth often differs in different datasets. The threshold suggested by the authors was derived from their dataset, thus may be only suitable for their data. If applied to other dataset, more training data and statistical models are required. Moreover, the authors should perform MeRIP-RT-qPCR or SCARLET experiments to validate that those low coverage peaks are not true m6A peaks.

It is important to clarify that we did not use a threshold to detect $m^6A$ peaks, and we have updated the methods to make this more obvious (line 486). We did use a 10-read or 10X filter *after* peak detection in two analyses to account for differences in sequencing depth and gene expression between experiments and conditions, as detailed below.

Sequencing depth is important for peak detection only to a point (as we show in **Supplementary Figure 2**). More directly, coverage in a particular gene or region determines whether a peak can be called. For **Figure 2**, comparing between data sets without establishing a threshold for gene expression would lead to underestimates of $m^6A$ peak concordance (e.g. if we included genes or exons that were expressed only in one data set but not the other). We therefore considered only peaks for genes expressed above a mean of 10X coverage to more fairly assess replicability of $m^6A$ detection while taking into account differences in gene expression and sequencing depth. We selected these thresholds based on **Figure 1a**, in which we analyzed two data sets: one our own and one from Engel et al., 2018. The higher threshold of 50X, where the Engel et al. data starts to plateau, proved too stringent for **Figure 2**, as there was insufficient gene overlap at this threshold for many pairs of data sets. To verify that our threshold of 10X was appropriate for other data sets, we include a new **Supplementary Figure 1c**, which summarizes across all experiments in **Figures 1-2**. This new figure shows that a mean gene coverage of 10X represents a reasonable approximation of the coverage necessary to detect most peaks across data sets.

To further measure peak detection accuracy, we generated additional MeRIP-RT-qPCR data to determine whether peaks below our threshold of 10 input reads are true $m^6A$ sites and found that 5/7 of the sites we tested showed METTL3-dependence, compared to 6/8 for peaks above our threshold (new **Supplementary Figure 1e**). Recent data from a new method to detect $m^6A$ using endoribonuclease digestion suggests that many $m^6A$ sites are missed by MeRIP-seq data (Garcia-Campos et al., 2019 (4)), thus it is perhaps not surprising that even low coverage peaks include true $m^6A$ sites. We have added clarification in the text regarding how the threshold should be interpreted: "These thresholds do not mean that peaks in genes with mean coverage < 10X or peaks with fewer than 10 input reads are false positives, but that the likelihood of false negatives rises with lower coverage (Additional File 2: Supplementary Figure 1e)." (lines 146-148).

We also include a threshold of ≥ 10 input reads per peak in **Figure 4a** across two conditions, where we were looking for changes in $m^6A$ and not just the presence of $m^6A$. We show the results without any threshold in coverage in **Supplementary Figure 4b**, which does not affect our conclusions. It does, however, illustrate that QNB in particular detected more peak changes where there was low expression in one or both conditions, which is of interest as it's unclear whether this is because these sites represent true changes or more noise at low expression levels.

3. In Figure 3a and Supplementary Figure 3b, the numbers of differential peaks detected by DESeq2, edgeR and QNB were very different from each other. For example, in siZc3h13 experiment, the edgeR identified tens of peaks, but DESeq2 and QNB predicted thousands of peaks. It seems to be the lack of standard algorithms for the detection of differential peaks. As such, using these programs to estimate the differential peaks may be inappropriate. In addition to the number of the union, the authors should also provide overlapping peak number among these three methods.

The lack of standard analysis methods in this field has clearly contributed to discordant reports on $m^6A$ dynamics. Our paper provides a starting place to compare methods and to understand the limitations of the underlying data. Of note, there is no ground truth data set in which the relative $m^6A$ enrichment at every site is known and validated under two conditions to assess differences in tool accuracy. As it is not possible to create such a data set using current methods, the best positive controls available are samples in which the methylation machinery has been disrupted, which is what we and others (Liu et al., 2017 and Cui et al., 2018 (5,6)) have used for tool evaluation, while the best negative control data sets are replicates at baseline conditions. However, we were unable to determine which of the three tools universally performed best on the positive/negative control data sets. Therefore, we suggest that in $m^6A$ peak analysis, researchers should consider starting with all three or using the intersects between 2 or 3 as a metric to rank peaks before follow-up experiments. We have now implemented an R package to facilitate this (https://github.com/al-mcintyre/deq). We have emphasized in the discussion and in the caption for **Figure 6** that additional validation of predicted changes is required. We have added the intersect to the results in **Figure 4a** and **Supplementary Figure 4b** (formerly Figure 3a and Supplementary Figure 3b), but it is not surprising that the overlap is poor.

4. The authors re-analyzed the previously published MeRIP-seq data but didn't provide the source codes and any running parameters of bioinformatics programs used. To guarantee the repeatability of their analysis, the authors are strongly recommended to upload the source codes to Github and list all running parameters and analysis pipelines.

In our manuscript, we cite the existing tools used and did upload our own code to Github, as described in the data availability section "Scripts used for analysis are available at https://github.com/al-mcintyre/merip_reanalysis_scripts" (lines 604-605). Many of those scripts depend on intermediate files too large to upload to GitHub (especially considering we reanalyzed data from 35 different studies); however, as noted in the response to Comment 3, we have therefore also implemented a pipeline in R to conveniently run the three tools we recommend and integrate results, available at https://github.com/al-mcintyre/deq.

5. The sensitivity and specificity are very important for bioinformatics analysis, which are, however, not adequately estimated in their analysis pipelines. In particular, the authors did not provide any positive control results to confirm the sensitivity and specificity of their analyses.

Here, it is important to distinguish sensitivity and specificity for peak detection and peak change detection.

In terms of peak detection, please see our responses above for additional validation of each peak caller with new MeRIP-RT-qPCR experiments with siMETTL3/14, which suggests that most predicted sites correspond to true positives.

In terms of peak change detection, we present the predictions for positive control data sets (with methylation machinery interference) and negative control data sets (replicates) in **Figure 3b**. However, as noted by Liu et al. (2017) (5) and above, there are no ground truth data sets in which the locations of $m^6A$ changes are well-characterized across the transcriptome, so calculating sensitivity and specificity is difficult, if not impossible. Cui et al. (2018) defined specificity and sensitivity using simulated data to suggest that MeTDiff is a highly accurate tool for peak change detection, but our study suggests that those results based on simulated data may be misleading (**Figure 3b-d**). In the biological positive controls we rely on instead, we do not expect every site detected as changed to reflect a true change in $m^6A$ for several reasons:

1) the direction of change in peak does not consistently match the expected direction of change for the interference (see figure **Response Figure 1** below for the number of peaks either increased or decreased with p adj < 0.05 and new **Supplementary Figure 3c** for the distributions of peak-gene $\log_2$ fold changes in our positive control data sets) – this was true for all of the methods tested and replicated using either MACS2 or MeTDiff for peak calling. It may be that as $m^6A$ is lost with Mettl14 knockdown (for instance), excess m6A antibody is redistributed to repetitive regions or other regions the antibody preferentially binds in the absence of $m^6A$ (see Lentini et al., 2018 paper on antibody biases, for example (7)), creating legitimate increases at peak sites that do not correspond to increases in $m^6A$.

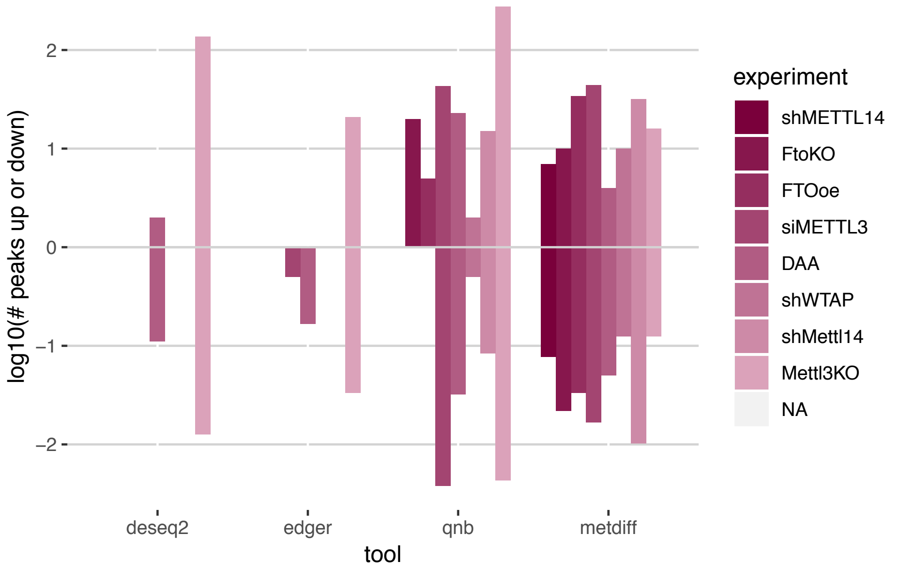


**Response Figure 1.** The number of peaks per tool that show either significant (adjusted p < 0.05) increases or decreases in response to various modes of interference with methylation machinery.

2) knockdown efficiency is variable and is known to affect results, and some methylation machinery knockdowns may not show any changes in $m^6A$ (see the comparisons of WTAP, METTL3, and METTL14 knockdowns in Schwartz et al., 2014 (8)), and
3) although we do already include an FTO experiment among our positive controls, there is still controversy over whether FTO is an active demethylase at $m^6A$ sites and the latest evidence indicates it is not (Garcia-Campos et al., 2019 (4)), suggesting that its utility as a control may be limited.

Because of the inherent ambiguity of the data, we cannot calculate sensitivity and specificity for these analyses.

6. Because m6A modification is tissue- or cell-specific, the positive and negative controls used for evaluating the performance of their approach in detecting m6A(m) peak changes should derived from same tissue or cell line. It's incorrect for the authors to use the negative control of datasets from mouse cortex and Huh7 cells in all conditions apparently with different tissues or cell lines.

Please let us clarify. Our evaluations of peak changes in **Figure 3** do not rely on cell type similarity between the positive and negative controls, as we have defined them. The positive controls consist of experiments featuring perturbation of the $m^6A$ machinery and the negative controls consist of identical biological replicates.

For each of our positive controls, we ran peak change detection between samples in which there was interference with methylation machinery (eg. Mettl3 knockout or shWTAP) and samples in which there was no such interference (eg. wildtype or shControl) from the same experiment. The latter could be considered internal negative controls, in which case they were from the same tissue or cell line, however, they were not the negative controls we used for comparisons of p-value distributions. Our negative controls for p-value distributions were the only two published data sets in which there were sufficient replicates to divide them into two groups for comparison under the same baseline condition. We have clarified this in the manuscript (line 225-228).

Under the null hypothesis of no differences in $m^6A$, a p-value distribution should be uniform. Therefore, regardless of cell type, we would expect that groups of replicates would show minimal differences in $m^6A$ between groups and more uniform p-value distributions than the positive controls. Interference with methylation machinery in the positive controls, meanwhile, should lead to shifts in $m^6A$ deposition and increases in detectable peak changes, and so we would expect to see leftward shifts in p-value distributions. As noted in our response to Comment 5, we expected variability in p-value shifts among positive control experiments, depending on the gene targeted and knockdown efficiency, which is why we included many experiments for this analysis. Our assumptions are supported by the new **Supplementary Figure 3c**, which shows that the peak-gene $\log_2$ fold changes for the negative controls center around zero, while those of the positive controls show show minor shifts that vary in magnitude and direction depending on the experiment.

We also note that while it is reasonable to hypothesize variation by cell type, work on this question is ongoing. Those who have suggested clustering of results by tissue based on MeRIP-seq have clustered based on IP reads, without accounting for differences in gene expression (e,g. Xiao et al., 2019 in Nat Cell Biol (9)) or have shown only "mild tissue-specificity" (e.g. the paper from Liu et al., 2019 recently accepted in Molecular Cell, for which the data will be released in 2020). The most compelling studies thus far are those that quantify $m^6A$ at particular positions to show differences. The original SCARLET paper found that at the locations they studied in *MALAT1*, the percent of transcripts methylated at a particular site could vary among cell lines, but the relative enrichment of $m^6A$ across sites was similar (see **Table 1** of Liu et al., 2013 (10)) – unfortunately they don't provide quantification of $m^6A$ machinery expression levels for comparison, but this could be a factor in the differences. A recent paper which used an endoribonuclease-based method for analyzing m6A sites genome-wide found that sequence context explained much of the variation of $m^6A$ levels across sites and suggested that "the primary mode through which $m^6A$ is likely to undergo modulation is through global regulation of $m^6A$ levels", for example through differences in the expression of methyltransferase complex components (see **Discussion** of Garcia-Campos et al., 2019). While MeRIP-seq cannot quantify $m^6A$, our results in **Figure 2a-b** show that technical or biological variation among experiments masks any variation in $m^6A$ detection by cell type. We have also added a new reanalysis of data from Xiao et al. (2019) (9), which shows little clustering by tissue type when considering peak overlaps for genes expressed with mean coverage ≥ 10 (as in **Figure 2a-b**), suggesting that identified $m^6A$ sites change little across cell type (see new **Figure 2c**) even if global $m^6A$ levels and the percent of transcripts methylated at individual sites do vary. (Clustering with a higher threshold of mean coverage ≥ 50 showed similar results, **Response Figure 2** below).

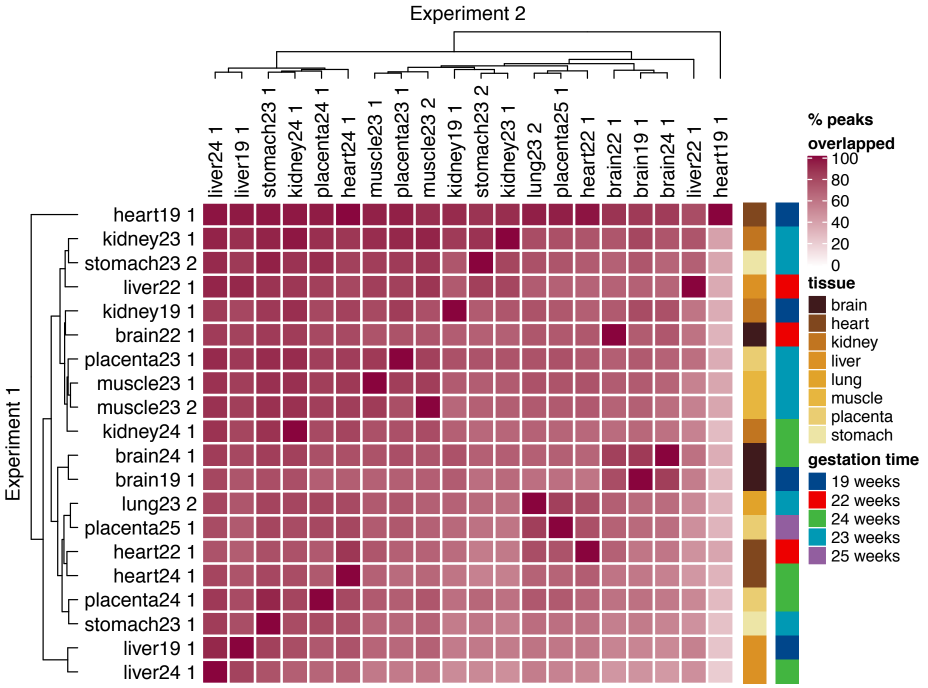


**Response Figure 2.** The percent of peaks detected in Experiment 1 also detected in Experiment 2 for genes expressed above a mean coverage threshold of 50 in both. Data from Xiao et al., 2019.

7. In Figure 3b, 3c, Supplementary Figure 3a,3e, there are multiple peaks with more than 2 fold-change between two conditions as displayed. It is highly suggested that y-axis of these figures should be normalized to same scales. Moreover, it is inappropriate for the authors to use the software without peer-review or confirmed by other biologists to draw coverage changes of MeRIP/m6A-seq.

One of our points, illustrated in **Figure 3d** (formerly Figure 2d), is that using the same y-axis scale masks differences in gene expression between two conditions or experiments. In analyzing MeRIP-seq data, it is not the total IP coverage that is important, but the coverage relative to the input RNA-seq experiment, and this input coverage changes as a function of gene expression. In order to see these changes, the same y-axis scales cannot be used, therefore we have decided not to modify **Figure 4** and **Supplementary Figure 4** (formerly Figure 3 and Supplementary Figure 3).

The code that we used is available on Github, as described in the manuscript: "Gene coverage was plotted using CovFuzze (https://github.com/al-mcintyre/CovFuzze), which summarizes mean and standard deviation in coverage across available replicates" (lines 559-560). A CovFuzze plot is a variation on a coverage plot, a standard type of representation for sequencing data. This method has been through peer-review before and was published as part of Imam et al., 2018 (11), to which we have now added a reference in the manuscript.

8. The authors declared that m6A peak overlapping in mRNAs from different studies is low. In fact, in Figure 1c, for HEK293T, HepG2 and mESC cell lines, the peak overlapping is relatively high. Moreover, studies from different groups all identified the standard m6A motif (RRACH) from their MeRIP/m6A-seq data, which well-justify the creditability of both the methods and results reported in different studies. It should be noted that the cell states (e.g. different stage of cell cycle), experimental conditions, and sequencing depth do influence the results leading to some different peaks in different studies. But this disparity among different studies objectively exists not only in MeRIP/m6A-seq but also in miCLIP, m6A-CLIP-seq, and ChIP-seq data. The authors should analyze the miCLIP and m6A-CLIP-seq data (also for m6A) and then compare the results with MeRIP/m6A-seq in order to estimate the reproducibility of these methods in different biological replicates and studies. For exploring the m6A change, it is highly recommended to first determine which experimental method is more suitable for this analysis.

The peak overlap reached a median of only 45% (within HEK293T 35%, HepG2 57%, MEF 25%, and mESC 53%). While not every data set does show enrichment of the $m^6A$ motif under peaks (see Ke et al., 2017 **Supplementary Figure 8** (3)), we agree with the reviewer that, indeed, most do show such enrichment. However, the question we wished to answer here was whether different experiments detect the same $m^6A$ locations or mutually exclusive subsets of $m^6A$ sites. Based on the overlaps of often <50%, we find that the situation is closer to the latter: MeRIP-seq experiments detect different subsets of $m^6A$ sites. We have added clarification to the text: "With rare exceptions (e.g. that described by Ke et al., 2017 in their Supplementary Figure 8 (3)), most MeRIP-seq data sets do show enrichment of the $m^6A$ motif DRAC. These results suggest, however, that multiple labs running MeRIP-seq on the same cell type will detect different subsets of $m^6A_{(m)}$ sites" (lines 184-187). We also appreciate the reviewer's clarity and have added the language suggested, "Possible contributing factors in the differences among studies include cell state (e.g. different stages of the cell cycle), experimental conditions, and sequencing depth." (lines 187-188). We note that ChIP-seq data presents a simpler problem for analysis, as it lacks the complexity added by differences in transcript abundance and isoform switching. Our analyses show that transcript expression variability is an important factor in analyzing MeRIP-seq data.

Although miCLIP was published in 2015, only three years after MeRIP-seq, the vast majority of $m^6A$ studies still use MeRIP-seq only (summarized in **Supplementary Table 1**). To our knowledge, this includes all sequencing data sets used to suggest changes in $m^6A$, with the exception of the heat shock data from Meyer the al., 2015 (shown in **Figure 4**), a data set from Zhang et al., 2018 on fear conditioning in mice, a data set looking at METTL3 knockdown from Vu et al., 2017, and the Ke et al., 2017 paper that examined differences among cell fractions but found no changes (3,12–14). There are several reasons why miCLIP is less widely used, including the complexity of the CLIP protocol and its lower sensitivity in the detection of $m^6A$ sites. Because our primary focus was testing statistical approaches for the detection of changes in $m^6A$ and reanalyzing the existing evidence for changes, we believe validation of a new analysis pipeline and re-analysis of the limited miCLIP data available is beyond the scope of the current paper.

9. In line 123-124, the authors provide erroneous conclusion of "m6A(m) presence does not decrease with expression level" that is contrast to all published papers. In fact, m6A promotes mRNA decay and directly reduces mRNA stability/expression level (Roundtree et al. Cell. 2017;169(7):1187-200.; Geula et al. Science. 2015;347(6225):1002-6.). This incorrect conclusion may derive from the arbitrary cutoff to filter the peaks in their analysis, which should be avoided.

We agree that, if anything, $m^6A$ increases at lower expression levels and have now added the additional Geula citation. Our point is that there is no known biological explanation for the decrease in $m^6A$ observed as transcript expression decreases (**Figure 1a**). Thus, we would expect this to be a technical artifact of the fact that peak calling is more difficult at lower expression levels. We have clarified the text to read: "Previous reports have suggested that $m^6A_{(m)}$ presence does not decrease with lower mRNA expression level, and, if anything, is higher in mRNAs with lower expression as methylated transcripts tend to be less stable (15,16). Peak callers, however, identify fewer peaks in genes at low expression, which we therefore assume reflects inadequate coverage for peak calling" (lines 133-137). **Supplementary Figure 4b** shows the results of our reanalysis without the threshold for input read counts, which does not affect our conclusions.

10. In Supplementary Figure 4, the authors identified m6A changes (13 positive correlations VS 6 negative correlations) when confirming MeRIP/m6A-seq by MeRIP-RT-qPCR, which is contrast to the conclusions in the whole manuscript. This issue further hints that the software used for calling peaks and identifying m6A changes in their analyses are inappropriate.

We are unclear on how the **Supplementary Figure 5** (previously Supplementary Figure 4) contrasts with the conclusions of the manuscript. We do not conclude that $m^6A$ changes cannot be identified, but that the data has technical limitations and considerations. We note in the discussion that our reanalysis of other papers suggests "meagre support for widespread changes in m$^6$A across the transcriptome independent of changes in the expression of methylation machinery."

**Supplementary Figure 5** expands on the comparison between MeRIP-seq and MeRIP-RT-qPCR shown in **Figure 5** for changes in m$^6$A predicted in response to viral infection. The correlations are calculated based on slight differences with infection by three species of *Flaviviridae*, all compared to uninfected cells. The six negative correlations are not surprising because for the changes we were able to detect in MeRIP-seq data, all three viruses produced similar directions of change, and we would expect that biological and technical variability could contribute to negative correlations when comparing small differences among viruses. When summarizing across genes in **Figure 5c**, the positive correlation between changes in enrichment detected by MeRIP-seq and MeRIP-RT-qPCR is clear. For further discussion of these data, see Gokhale et al., 2019 (bioRxiv), our paper recently accepted in Molecular Cell (17).

Minor comments:

1. The authors aligned m6A-seq reads to the genome instead of transcriptome, which will miss the reads spanning exon junctions and is incorrect for handling m6A-seq data.

We agree that accounting for splice junctions in RNA data is important. As the Reviewer notes in the next comment, we aligned using STAR, which is an aligner designed for RNA-seq data that takes into account splice junctions. We have added this description to the methods (line 478): "STAR, a splice-aware aligner for RNA-seq data." For further information, please see https://doi.org/10.1093/bioinformatics/bts635 (18).

2. The authors should provide the parameters used for STAR aligner.

We have added a description of the non-default parameters used and another link to the GitHub page with analysis scripts.

3. Given FTO is an m6A demethylase, the authors should analyze the FTO-KO data (GSE47216) in their pipelines and display the results in Fig 3a and Supplementary Figure 3b.

We have now included this additional study, although as we note in the text, there is mixed evidence that FTO is an m$^6$A demethylase (see also the recently published paper from Garcia-Campos et al., 2019 (4)).

4. Ke et al. [Genes Dev 2017, 31(10):990-1006)] employed m6A-CLIP to identify m6A change in Mettl3 KO mES cells by calling peaks. The authors should use their pipeline to analyze the data (GSE86336) and display the results in Fig 3a and Supplementary Figure 3b. This data will help the authors to evaluate their pipelines and meanwhile enable biologists to compare the accuracy between m6A-CLIP and MeRIP/m6A-seq data.

The benefit of miCLIP/m$^6$A-CLIP, which follows a protocol similar to MeRIP-seq in terms of antibody enrichment of methylated fragments but introduces greater technical challenges with crosslinking, is increased resolution for the localization of m$^6$A sites to single bases. Most analyses of miCLIP/m$^6$A-CLIP data involve the detection of crosslinking-induced mutations or truncations, rather than peaks alone. We therefore suggest that very different tools would be more appropriate for this type of analysis. We would further expect that, as the reviewer notes in their Comment 8, "disparity among different studies objectively exists not only in MeRIP/m6A-seq but also in miCLIP, m6A-CLIP-seq and ChIP-seq data." Using a single miCLIP data set for comparison could skew expectations, depending on the quality of that particular data set, and whether it was representative. All in all, miCLIP data requires further study, and we have updated the discussion to refer to Garcia-Campos et al., 2019, a paper that makes several interesting observations in the comparison of this technique to an endoribonuclease digestion-based method for the analysis of m$^6$A sites (4): "So far, comparison of this data to miCLIP suggests that despite poor overlap among miCLIP studies, most sites identified by miCLIP are true m$^6$A sites, and that higher m$^6$A:A ratios are associated with identification of a site in more studies2020-01-09 4:55:00 PM. However, Garcia-Campos et al. also suggest that antibody-based approaches may underestimate the number of m$^6$A sites" (lines 427-431).

5. The authors did not provide any information about how they calculate the read counts for each peak. The tools used to calculate read counts for each peak should be described.

We have added this description: "Reads aligned to peaks were counted using featureCounts from the Rsubread package (19)" (line 499). All of our code used in this analysis is also available on the DEQ GitHub page.

6. The authors only provided figures to display the number of peaks but didn't list the detailed peak information from different studies, which makes it difficult to reproduce and check the results. All peaks (BED12 or BED6 format) should be listed.

We have added an **Additional File 4** with the output files from DEQ, including the peak locations and p-values from DESeq2, edgeR, and QNB.

7. Different MeRIP/m6A-seq experiments were chemically fragmented into tags with different length. However, the authors did not provide any detailed information (e.g. shift size for each tag) when they call peaks using MACS and identify m6A changes using DESeq2, edgeR and QNB software. Moreover, the authors should use the length of fragment to calculate the coverage or read counts of each peaks/genes.

Size shifts are not provided for many published studies; therefore, in some cases, we are not able to report that information. However, where estimated fragment lengths were reported (generally approximate means) or could be estimated from paired end data, we have added them to **Supplementary Tables 2-5**. For studies in which we were unable to find fragment length information, we estimated fragment length for the purposes of peak calling and read counting based on the median across studies (100 bases).

8. Detailed information about the methods and parameters used in the manuscript remains totally unclear. A whole pipeline should be provided and all source codes should be uploaded to Github. Moreover, all methods and parameters/ cutoff in the published studies should be listed and what parameters/cutoff is important should be discussed in this study.

The scripts are all uploaded to GitHub and publicly available, as noted above and as linked in the appropriate section of the text ("Availability of Data and Materials"). We have also updated our materials and methods section, and further summarize the information available on analysis in the methods sections of other studies in **Supplementary Table 1**.

**Reviewer #2:**

N6-methyladenosine is the most abundant internal mRNA modification. In 2012, two groups independently developed MeRIP-seq/m6A-seq and first mapped m6A methylome in the transcriptome wide manner. Since then, this method has been widely used to map m6A in different species, biological processes and stress conditions. And many studies have shown that m6A is dynamic regulated and plays important and diverse roles in these biological processes. In this study, the authors reanalyzed these published datasets and compared the tools used for statistical analysis by these studies and they claimed that fewer changes can be detected compared to the original reported sites and the detection reproducibility is limited between different studies. The problem of low reproducibility between different studies indeed occur in those technologies dependent on immunoprecipitation, however the authors just compared the existing statistical analysis tools and do not provide an effective solution to solve it. More data and bioinformatic analysis are needed to support their conclusions.

We thank the reviewer for their comments noting the issues with technologies dependent on immunoprecipitation. These issues have not been deeply explored before for MeRIP-seq, as evidenced by the continued publication of studies that use few replicates and no or inappropriate statistical tests to draw conclusions that are poorly supported by the data and irreproducible between studies. To help with this, we have used many of the MeRIP-seq data sets published to date and created an open-source computational tool to conveniently run comparisons of peak enrichment (DEQ). Our overall pipeline for differential peak calling outlined in **Figure 6** can provide guidance for future MeRIP-seq analyses.

Specifically,

1. To evaluate the changes of differentially methylated transcripts under stress conditions, the authors picked several transcripts and plotted the coverage of Input and IP under different conditions, and made a statement that the reproducibility of these studies is poor. However, it is not a proper way to evaluate the reproducibility of these methylation changes by picking several sites and plotting the coverages, as false positive sites exist in all high throughput sequencing methods. The authors should provide the false positive rate of these datasets, not just show some examples.

Strikingly, we were unable to detect reproducible peak changes in any of these data sets except for four sites noted in the dsDNA response experiments, and we have added clarification to that effect: "Applying the same statistical approaches, we were likewise [similar to the KSHV comparison] unable to detect any shared peak changes between the studies of HIV infection, and there were insufficient replicates to compare heat shock studies (20–24). Thus, in our reanalysis of $m^6A$ changes in response to stimuli, we detected only four statistically reproducible peak changes, all in response to dsDNA" (lines 330-334).

Also, please see our response to Comments 3 and 5 from Reviewer #1. Unfortunately, we can't calculate a false positive rate for a data set without knowing the ground truth (the differential $m^6A$-status of any particular RNA site). Failure to replicate a peak change does not necessarily mean it is a false positive, as it could also indicate insufficient power or some other unanticipated source of variation (e.g. quality of the antibody or immunoprecipitation).

2. To detect methylation changes between conditions, the authors compared several tools used for statistical analysis and found that tools that account for overdispersion are better. However, there are no improvements in this study, as these tools already exist. The authors should provide a new pipeline or optimize the parameters of these statistical analysis tools to robustly detect the methylation changes.

The generalized linear model approaches we describe had not previously been applied to MeRIP-seq data. We show here that they are reasonable methods for this analysis based on our evaluations of positive and negative control data sets. In general, we hope that our evaluations will prompt biologists to adopt better validated methods, as there is no consensus in the field on methods to analyze MeRIP-seq data. To facilitate this, we have implemented a pipeline in R to conveniently run the three tools we recommend and have added a link under "Availability of Data and Materials": https://github.com/al-mcintyre/deq.

3. The authors suggested that 6-9 replicates are needed to detect the consistent peak changes under different conditions. However, it is difficult to perform so many replicates, especially for these samples difficult to obtain. In addition, as the authors stated, more replicates can only increase the peak numbers but not the DRAC motif enrichment, indicating that more false positive sites will be included. Hence increasing the replicates number is not a good solution to ensure the detection consistence.

We have updated the manuscript to clarify a difference here in terms of peak detection vs. peak change detection. The benefits of increased replicates to peak detection are, as the reviewer notes, indeed not related to motif enrichment, but **Figure 5** shows the benefits to peak change detection. We appreciate that, especially for clinical samples, it is inconvenient to get so many replicates, but those who are planning experiments should be aware that only a small subset of peak changes may be detectable with fewer replicates. We have added a comparison to recommendations for RNA-seq studies, which, though the experiments are simpler, are in line with our own:

"Schurch et al. (2016) and Conesa et al. (2016) produced similar recommendations for basic RNA-seq studies, finding that 6-12 replicates were necessary to detect most changes in gene expression and that changes of 1.25 were detectable 25% of the time with five replicates, rising to 44% with ten replicates, respectively. While our results broadly agree with these recommendations for RNA-seq, they also suggest that almost all published MeRIP-seq studies to date are underpowered" (lines 367-372).

4. They analyzed the overlap of peaks among studies and claimed that peaks showed higher overlap within different cell types from the same study than within the same cell type from different studies. Conceptually, it makes no sense to compare the detected peaks among different

studies, as the choice of antibodies and IP conditions have a huge impact on peak detections. Therefore, researchers compared the samples in the same batch to identify the differentially methylated transcripts. In addition, the culture conditions and origins of the same cell line are different in different labs, which can influence the m6A methylome of the cells. Hence, the difference of the detected m6A peaks on the same cell type by different labs also indicates the m6A is dynamic regulated under different conditions.

We checked for experimental factors that could have contributed to the differences in peak detection within cell types (see **Supplementary Table 2** and lines 181-183), however we were not able to find a clear correlation with antibody choice or experimental protocols. Of course, we do not know the exact culture conditions and cell origins for each of the studies and cannot rule out these or other associated factors. However, we think that for any experiment in biology, the default assumption should be that the data will be replicable in other labs, and if it is not, the extent to which it varies is worth pointing out. Comparing experiments from different batches and labs can be confounded by many variables, but as discussed in our response to Reviewer 1, **Figure 2a-b** shows the extent to which $m^6A$ detection differs among experiments. Without further experiments, it is impossible to determine whether differences in detected $m^6A$ sites among studies are due to dynamic regulation under different conditions. Interestingly, tissues from the same study (Xiao et al., 2019) showed high peak overlap even though samples were taken from different fetuses and time points (new **Figure 2c**), suggesting that sample processing is a large component of the variation, rather than biological dynamics (9).

5. The authors used the notion of variance explained incorrectly. It is R2 (variance) instead of R (correlation coefficient) that represent the variance explained.

In our text, we did not use the term "variance explained." We do take correlations and report the correlation coefficients, however, we don't suggest that R represents the variance explained, and we have ensured this is clear in the updated manuscript as well. We appreciate the need for clarity on this point.

Minor points,
1. The authors stated that "the percent of peaks detected in one experiment that were also detected in a second varied among pairs of studies from as low as 2% of peaks to as high as 90%". To describe the variance among studies, the median value should be shown not the maximal and minimal rate.

While the median was previously included in the discussion, we have now added it to this section of the results as well: "as low as 2% of peaks to as high as 90% (median = 45%)" (line 178), as suggested by the reviewer. We agree this is helpful for context of the peak overlaps.

2. The authors used MeRIP-RT-qPCR assay to validate the peaks detected by MeRIP-seq. However, this method also relies on antibody immunoprecipitation and it can only exclude the possibilities of sequencing error or duplications. Hence, the orthogonal methods are still needed for validation.

Validation, in addition to reproducibility, remains an issue for the field in general. Unfortunately, methods available for validation are limited. Although we tried SCARLET for four sites, the protocol did not work for our sites of interest, either because the expression of these genes is too low (below the recommended FPKM of > 50) or because the oligos were not specific enough. A new endoribonuclease method for detecting $m^6A$ (Garcia-Campos et al., 2019 (4)) works only for ACA sites far enough from other ACA sites for amplification or sequencing (~16% of sites).

However, we note that in the revised manuscript, we provide additional validation of peak detection to compare between peak-callers. We tested peaks detected by MeRIP-seq using MeRIP-RT-qPCR on RNA from cells in which the $m^6A$ methyltransferases METTL3 and METTL14 were depleted (**Supplementary Fig 1b**). We found that 4/4 peaks called by MACS2 and a majority of those called by MeTPeak and MeTDiff showed reduced $m^6A$ enrichment in METTL3/14 depleted cells, which indicates that our MeRIP-seq analysis detects peaks that are dependent on METTL3/14 and likely true modifications.

### Response references

10. Liu N, Parisien M, Dai Q, Zheng G, He C, Pan T. Probing N6-methyladenosine RNA modification status at single nucleotide resolution in mRNA and long noncoding RNA. Rna. 2013;

**Final responses received:**

Reviewer #1: The authors have tried to improve this manuscript, but they do not show advanced techniques and new experimental evidence to answer the existing question.

Reviewer #2: In this revised version of their manuscript, the authors have performed some additional analysis and added some information to address the reviewer's concerns. Indeed, the authors have raised a new pipeline for m6A peak calling, however the novelty of this study is limited as they used the existing analysis methods and just optimized the parameters. In addition, as the authors mentioned in the new Figure 2c, the experimental factors are dominant factors leading to the low overlap rate and optimization of bioinformatics analysis pipeline cannot effectively increase the reproducibility between different studies. Hence, the choice of bioinformatics pipeline is less important compared to the differences from experimental factors. Besides, using 6-9 replicates for one sample is difficult to achieve for most studies. Overall, I wonder whether the new pipeline can benefit future studies and this study may not meet the criteria of Genome Biology.
